# Nonredundant roles of the HDAC3 corepressor complex subunits SMRT and NCOR in controlling inflammatory and metabolic macrophage pathways

**DOI:** 10.1101/2025.05.13.653058

**Authors:** Astradeni Efthymiadou, Chaode Gu, Cheng Wang, Hongwei Wang, Ziyi Li, Oihane Garcia-Irigoyen, Rongrong Fan, Eckardt Treuter, Zhiqiang Huang

## Abstract

Transcription factors and coregulators coordinate inflammatory and metabolic pathways in macrophages through epigenetic and transcriptional mechanisms. The HDAC3 corepressor complex plays fundamental roles in these mechanisms, with the homologous subunits SMRT and NCOR being critical for complex assembly and interactions with transcription factors and chromatin. However, the relative contribution of SMRT and NCOR in controlling complex-dependent macrophage pathways remains poorly understood. Here, we assessed their genome-wide roles in mouse macrophage RAW264.7 cells and in bone marrow-derived macrophages. Transcriptome analysis upon corepressor depletion identified six differentially expressed gene clusters. SMRT depletion primarily upregulated inflammation-related pathways, NCOR depletion primarily upregulated metabolism-related pathways. Epigenome analysis revealed that corepressor depletion differentially altered chromatin accessibility and H3K27 acetylation, consistent with transcriptome changes. Cistrome analysis revealed that both corepressors differentially influence each other at chromatin. SMRT uniquely controls the chromatin binding and nuclear localization of NCOR, GPS2 and HDAC3, thus acting as the chromatin anchor for the corepressor complex. Finally, corepressor depletion differentially modulated macrophage reprogramming in response to TLR4, IL4 and LXR signaling. Overall, our study reveals a hitherto underappreciated non-redundant role of SMRT and NCOR in coordinating chromatin accessibility, H3K27 acetylation, enhancer activity and transcription to differentially regulate inflammatory and metabolic macrophage pathways.

**GRAPHICAL ABSTRACT:** 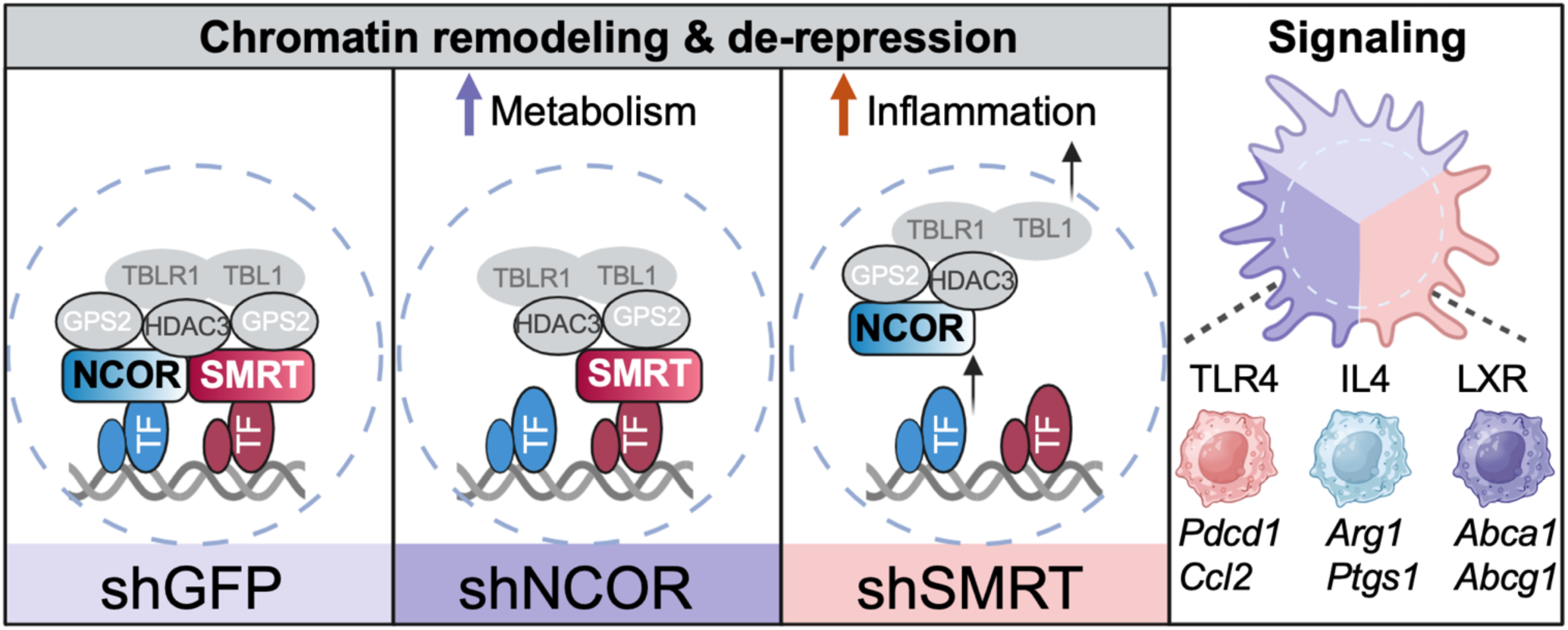

## INTRODUCTION

Macrophages are an integral part of the innate immune system. They are phenotypically diverse and even though their most widely known functions are the phagocytosis and degradation of pathogens and the activation of the adaptive immune system, they also have roles in development, tissue regeneration and lipid metabolism (1–6). Macrophages have been implicated in chronic inflammatory and metabolic diseases, such as type 2 diabetes, cardiovascular diseases, rheumatoid arthritis and cancer, making them attractive targets for therapeutic intervention (1,7–10).

Macrophage heterogeneity is attributed to distinct activation states, extending beyond the classical pro-inflammatory M1 and anti-inflammatory M2 states, in response to signals from the microenvironment (3,4,11–13). Macrophage activation is primarily regulated at the level of transcription and is characterized by the dynamic interplay between transcriptional activation and repression, which determines gene expression patterns linked to metabolic and inflammatory pathways (14,15). Via direct binding to cis-regulatory elements (i.e. enhancers, silencers, promoters), lineage-determining transcription factors (TFs), such as PU.1 and C/EBPs, and signal-responsive TFs, such as AP1 (e.g. JUN, JUNB, ATFs), NF-κB (e.g. p65/RELA), STATs, IRFs, nuclear receptors (e.g. LXRs, PPARγ), regulate gene expression to determine macrophage activation states (14). In addition to TFs, the synergy between signal transduction and epigenetic regulation via chromatin structure, histone modifications, and DNA methylation, plays a central role in gene expression, cellular responses and memory mechanisms in macrophages (11,14,16–20). Therefore, epigenetic alterations and disruption of metabolic and inflammatory signaling are closely associated with abnormal tissue function in disease, underscoring the significant impact of macrophage signaling and epigenetics on disease pathogenesis.

The signaling-dependent epigenetic regulation of gene expression critically depends on transcriptional coregulators (also referred to as ‘cofactors’), which interact with TFs and operate in multi-protein complexes carrying chromatin-modifying activities, thereby adding an extra level of complexity to the activity regulation of cis-regulatory elements, including promoters, enhancers and silencers (14,15,21–24). While coactivators, including CBP/p300 (the TF-binding H3K27 acetyltransferase implicated in multiple transcriptional activation steps) and MED1 (TF-binding core subunit of the mediator complex) are established regulators of enhancer function, corepressors have only recently been recognized to also specify enhancer function, which mechanistically may include direct coactivator antagonism (17,18,25).

The HDAC3 corepressor complex (also referred to as the ‘nuclear receptor corepressor complex’), is amongst the best-studied corepressor complexes and has particularly been implicated in the regulation of metabolic and inflammatory pathways (15,26–34). The complex includes the TF-interacting core subunits nuclear receptor corepressor (NCOR, alias N-CoR, NCOR1, encoded by gene *Ncor1*) (31,32), silencing mediator of retinoic acid and thyroid hormone receptors (SMRT, alias NCOR2, encoded by gene *Ncor2*) (33), and G protein pathway suppressor 2 (GPS2), further histone deacetylase 3 (HDAC3) and two transducing β1-like proteins (TBL1 and TBLR1). There is evidence that functionally distinct subcomplexes or modules operate in macrophages and in hepatocytes, despite all core subunits are expressed (15,27–29,35). However, the exact mechanism behind the formation and regulation of these subcomplexes is not yet understood. SMRT and NCOR are large homologous 270 kDa proteins with a conserved domain structure and biochemical features implicated in corepressor complex assembly, HDAC3 activation, and binding to many different TFs and to histones. However, various studies, including corepressor-depleted mouse and cell models along with mechanistic structure-function studies, suggest that SMRT and NCOR serve non-redundant functions (15,26,28). For example, phenotypes of macrophage-specific NCOR and HDAC3 knockout mice share similarities but seem distinct from the phenotype of GPS2 knockout mice. While macrophage-specific SMRT knockout mice have not yet been reported, henotypes of SMRT mutant (nuclear receptor-interaction domain deficient) mice suggest unique roles not shared with NCOR (36–41). Overall, it remains poorly understood which gene expression networks and pathways depend on each corepressor complex subunit, and whether SMRT and NCOR mechanistically operate through distinct mechanisms to specify the functions of the corepressor complex and its postulated subcomplexes.

Here, we address this open issue using a comparative genome-wide analysis of the SMRT vs. NCOR depletion-regulated alterations of transcriptome, epigenome and cistrome in the mouse macrophage RAW264.7 cell line. We additionally validate transcriptome and mechanistic data in mouse bone marrow-derived macrophages (BMDMs). Our comparative analysis identifies distinct roles of SMRT and NCOR, and their respective subcomplexes, in differentially coordinating chromatin remodeling, histone acetylation, and transcription to regulate signal-dependent macrophage activation states linked to inflammation, metabolism and cancer. These differences are indicative of distinct functional subcomplexes and altered preferences to target TFs and to chromatin. Finally, our study identifies a hitherto unrecognized functional hierarchy with SMRT but not NCOR controlling the nuclear localization and the chromatin binding of the remaining corepressor subcomplex, and thereby macrophage gene expression programs.

## MATERIALS AND METHODS

### Cell culture and treatments of RAW264.7 cells and BMDMs

The RAW264.7 mouse macrophage cell line, hereafter referred to as *‘*RAW cells*’*, was purchased from the American Type Culture Collection (ATCC, TIB-71) and cultured in DMEM with 10% heat-inactivated FBS (Gibco™, A3160802) and 100 U/mL pen/strep (Gibco™, 15140122), with incubation at 37°C and 5% CO2. RAW cells we used for analysis between passage numbers 10 to 25. For pathway activation, RAW cells were incubated for 6 hours incubation with 10 ng/mL LPS (SigmaAldrich, L4516), 20 ng/mL IL4 (Sigma, SRP3211), or 5 μM GW3965 (Sigma, G6295) followed by RNA extraction for RT-qPCR analysis or RNA-seq. The same treatments were applied for a 3-hour incubation for H3K27ac CUT&Tag in cells depleted of NCOR and SMRT. BMDMs were isolated from wild-type C57BL/6 mice (GemPharmatech, N000013) as previously described (37) and treated with lentivirus-shRNAs as outlined below.

### Lentivirus shRNA-mediated knockdown in RAW cells and in BMDMs

We designed lentiviral shRNA sequences targeting NCOR (*Ncor1*) (clone IDs: shNCOR-1 TRCN0000350169, shNCOR-2 TRCN0000310755) and SMRT (*Ncor2*) (clone IDs: shSMRT-1 TRCN0000238140, shSMRT-2 TRCN0000238139) using the GPP Web Portal (Broad Institute). These sequences were synthesized and constructed into the PLKO.1-TRC vector (Addgene, 10878). The lentiviral particles were then packaged in HEK293FT cells using psPAX2 (Addgene, 12260) and pMD2.G (Addgene, 12259). Subsequently, the lentivirus particles were transduced into RAW macrophages at passage numbers lower than 10, and stable cell lines were established through puromycin selection (5 μg/mL) over a 5-day period. The efficiency of the knockdown was tested at the mRNA expression level with RT-qPCR analysis and the protein expression level with Western blot analysis, see also our previous study (25). We evaluated two knockout cell pools for depletion of NCOR and SMRT each and selected one with best knockdown efficiency. Additionally, we knocked down NCOR and SMRT in BMDMs using the same lentivirus shRNAs (TRCN0000238140 for SMRT and TRCN0000350169 for NCOR) and conducted transcriptome RNA-seq sequencing. The shRNA sequences targeting GFP, NCOR and SMRT are provided in **Supplementary Table S2**.

### siRNA-mediated knockdown in RAW cells

RAW cells were transfected with control siRNA (Dharmacon, D-001810-10-20) or NCOR siRNA (Dharmacon, *Ncor1*, L-058556-00-0020,) or SMRT siRNA (Dharmacon, *Ncor2*, L-045364-00-0020). In brief, 5 × 10^5 cells were seeded per well in a 6-well plate and incubated overnight at 37°C with 5% CO_2_ to allow cell adhesion. The next day, 25 nM of the corresponding siRNA and 7.5 μL Lipofectamine RNAiMAX reagent were each diluted in 250 μL of OPTI-MEM medium. After a 10-minute incubation, the siRNA-Lipofectamine RNAiMAX mixture was added to cells in 1.75 mL of medium without antibiotics. After 6 hours, the transfection medium was replaced with complete DMEM containing antibiotics. The cells were then incubated for an additional 42 hours (48 hours post-transfection) before treatment with 10 ng/mL LPS for 6 hours. Subsequent RNA extraction, qPCR validation and RNA-seq were performed. The siRNA sequences are provided in **Supplementary Table S2**.

### RNA isolation and RT-qPCR

Total RNA was extracted from the cells (2×10^5^) using the E.Z.N.A. Total RNA Kit I (Omega, R6834-02), according to the manufacturer’s instructions. cDNA synthesis was performed with M-MLV Reverse Transcriptase (Life Technologies, 28025-021). RT-qPCR was conducted using SYBR Green master mix (KAPA BIOSYSTEMS, KK4617) and all samples were normalized to *Gapdh* mRNA. The primer sequences are detailed in **Supplementary Table S1**. Relative changes in mRNA expression were calculated using the comparative cycle method (2^-ΔΔCt^).

### RNA-seq

RNA was extracted as previously described and was sent to Novogene. The RNA quality was assessed using TapeStation (Agilent 4200) by the RNA Integrity Number (RIN). RNA-seq samples were sent to Novogene (Danmark) for library preparation and sequencing. Sequencing was performed on the NovaSeq 6000 platform (Illumina) with the PE150 parameter. Subsequently, the FASTQ data were aligned to the mouse reference genome mm10 using HISAT2 (42). The primary analysis of the RNA-seq data was executed in R 4.2.1. Z-normalized score values were utilized for principal component analysis (PCA) using the prcomp function from the base R library. Differential gene expression analysis was performed using DESEq2 v1.36.0 (43), considering genes with *P* adj ≤ 0.001 as significant. Gene expression heatmaps were generated using pheatmap v1.0.12 (https://cran.r-project.org/web/packages/pheatmap/index.html). Over-representation analysis using the KEGG pathway database was executed with clusterProfiler v4.4.4 (44) and the enriched terms were visualized as a network. TF enrichment analysis was conducted using ChEA3 (45), with the MeanRank output used as the ChEA3-score for downstream analysis. Enriched TF networks were generated with ChEA3 local networks.

### ATAC-seq

Assay for Transposase-Accessible Chromatin using sequencing (ATAC-seq) on cell nuclei was performed according to established protocols (46,47). NCOR and SMRT stable knockdown cells (50000 cells) were harvested (basal condition) and suspended in lysis buffer. and their nuclei were isolated. Cell nuclei were spun down and underwent a transposition reaction at 37°C for 30 minutes. Genomic DNA was extracted using the PCR Purification Kit (Qiagen, 28106). The ATAC-seq library amplification process was adhered to protocols outlined in our previous publications [26, 76]. The resulting purified DNA library mix was then sequenced on the NextSeq 550 platform (Illumina) (BEA Core Facility, Karolinska Institutet, Sweden) with pair-ended output, generating 75 pair-end reads.

### ChIP-seq and CUT&Tag

Chromatin immunoprecipitation followed by sequencing (ChIP-seq) was conducted according to our established methodologies (17,25,37). The antibodies used included H3K27ac (Abcam, ab4729), GPS2 (custom-made, as previously described) NCOR (Bethyl, A301-145A), and SMRT (Bethyl, A301-147A). In brief, RAW cells (shGFP, shNCOR, and shSMRT cells) from a 15 cm^2^ culture dishes were crosslinked with 10 mL of 2 mM disuccinimidyl glutarate (DSG) (VWR, A7822.0001) for 30 minutes, followed by 10 mL of 1% formaldehyde for 10 minutes. The crosslinking reaction was quenched by adding 500 μL of 2.5 M glycine to a final concentration of 0.125 M and incubating for 5 minutes. Subsequently, the lysed RAW cell nuclei were sonicated for 30 minutes (30s ON/30s OFF) using a Bioruptor Pico (Diagenode, B01060010). Protein A Dynabeads (Invitrogen, 10002D) were incubated with the specified antibodies (1-4 μg). The ChIP-enriched DNA was purified using the Clean & Concentrator Capped Zymo-Spin I kit (Zymo Research, D4013). NCOR, GPS2, and SMRT ChIP-seq were performed in both NCOR and SMRT depletion cells along with H3K27ac ChIP-seq. We repeated two independent NCOR ChIP-seq in SMRT depletion cells to draw robust conclusions. ChIP-seq libraries were prepared using the Takara ThruPLEX DNA-Seq Kit (Takara, R400736). Sequencing was performed on the Illumina NextSeq 550 platform (Illumina) with 75bp pair-end reads. CUT&Tag experiments were conducted following a published protocol (48) with the H3K27ac antibody. Knockdown cells were treated with LPS, IL4, or GW3965 for 3 hours, and these cells were subsequently used for CUT&Tag assays. Library samples were sequenced on the NextSeq 2000 platform (PE100) with paired-end output (BEA Core Facility, Karolinska Institutet, Sweden).

### Computational analysis of ChIP-seq, CUT&Tag, and ATAC-seq data

To acquire TF binding profiles in RAW cells and BMDMs, we analyzed data from our previously published ChIP-seq experiments (17,25) and relevant datasets from other GEO databases (**Supplementary Table S3)**. ChIP-seq data were extracted from the GSE184884, GSE50944, GSE106706 and GSE130383 series, encompassing GSM7501916, GSM7501917 (GPS2); GSM4848528, GSM4848529 (PU.1); GSM4848610, GSM4848611 (CBP); GSM7903391, GSM7903392, GSM7903393, GSM7903394, GSM4848503, GSM4848504, GSM4848507, GSM4848508 (H3K27ac); GSM1232940, GSM123294 (H4K5ac); GSM7903399, GSM7903400, GSM7903401, GSM7903402 (NCOR); GSM4848572, GSM4848573 (MED1); GSM4848548, GSM4848549 (RUNX1) and GSM4848584, GSM4848585 (JUNB). Sequencing files (FASTQ files) were aligned to the NCBI37/mm10 version of the mouse reference genome using Bowtie2 on the GalaxyEurope platform. Subsequently, the sequencing tags were processed and imported into HOMER (49). Peak identification was carried out utilizing HOMER with default settings, although slight variations in peak calling parameters were observed between ChIP-seq experiments for histone marks and TFs/coregulators. Overlapping peaks were ascertained by merging of individual peak files from each experiment. NCOR and SMRT ranked super-enhancers were done using the Rank Ordering of Super-Enhancers (ROSE) algorithm (50). For CUT&Tag data, MACS2 was employed for peak calling (51). Differential peak tag count analysis was performed using edgeR in R (52) with peaks demonstrating an adjusted *p*-value < 0.05 deemed differentially enriched. Bedtools was used to identify overlapping peaks for specific analyses and extract desired peak coordinates (53). Peak coverage analysis was conducted utilizing annotatePeaks.pl, focusing on the 3 kb upstream and 3 kb downstream regions from the indicated peak center. Heatmap plots were generated through the utilization of Deeptools (54). The correlation between the associated TFs and coregulators was assessed using the plotCorrelation function in Deeptools (54). In the case of ATAC-seq results, paired-end data were aligned to the mouse mm10 genome via Bowtie2, and ATAC-seq peak calling was executed using MACS2. Enriched TF motif sequences in the ATAC-seq, ChIP-seq and CUT&Tag peaks were identified using the motif enrichment analysis package by HOMER (49). Statistical analyses for differential expression were performed using edgeR, with peaks displaying changes and an adjusted *p*-value < 0.05 considered differentially enriched.

### Western blot analysis

Immunoblotting procedures followed our previously established protocols [26]. For each experiment, 2×10^5^ cells of NCOR, SMRT, and GFP knockdown RAW cells were seeded one day in advance and washed twice with PBS before cell lysis. Protein concentrations were determined using the BCA assay (Life Technologies, 23235). Subsequently, 20 μg of total protein samples were loaded onto SDS-PAGE gels (Invitrogen, NP0335BOX), transferred onto PVDF membranes (Cytiva, GE10600069), and subjected to antibody probing. Antibodies against NCOR (Bethyl, A301-145A, 1:3000), SMRT (Bethyl, A301-147A, 1:3000), HDAC3 (Beyotime, AF2011, 1:2000), GPS2 (home-made rabbit polyclonal, as described in (55), STAT6 (Sigma, S-6433, 1:3000), p65 (Abcam, ab32536, 1:3000), JUN (Santa Cruz, sc-45,1:3000), β-actin (Abcam, ab8226, 1:5000), Lamin B1 (Beyotime, AF1408,1:3000), GAPDH (Proteintech, 60004-1-Ig, 1:5000). The anti-mouse (EMD Millipore, AP308P, 1:5000) and anti-rabbit (GE Healthcare, NA934V, 1:5000) secondary antibodies were used accordingly. Membranes were incubated using ECL substrate (BIO-RAD, 170-5061) and revealed on Amersham Hyperfilm ECL (Cytiva, 28906836). Antibodies are listed also in **Supplementary Table S2**.

### Immunofluorescence (IF) sub-cellular localization analysis using confocal microscopy

We used two different lentivirus-shRNAs (shSMRT-1. shSMRT-2, shNCOR-1, shNCOR-2) to knockdown each corepressor in RAW cells. Cells were fixed in 4% paraformaldehyde for 30 minutes, permeabilized with 0.1% Triton X-100 in PBS, and incubated with anti-NCOR (Bethyl, A301-145A, 1:200) and anti-SMRT (Bethyl, A301-147A, 1:200) antibodies in 0.5% BSA/PBST at 4°C overnight. The cells were then incubated with fluorescence-conjugated anti-rabbit secondary antibodies (Beyotime, A0423, 1:1000) and DAPI (Beyotime, P0131, 1:1000) for 1 hour at room temperature. An FV3000 laser scanning confocal microscope (Olympus) was used for examination and image capture. According to our previous protocols (37) bone marrow cells were isolated from C57BL/6 mice and differentiated into bone marrow-derived macrophages (BMDMs) using GM-CSF (20 ng/mL). BMDMs were infected on day 7 of differentiation with lentivirus (shGFP, shNCOR-1 and shSMRT-1) for 24 hours, followed by selection with puromycin (5 μg/mL) for 7 days.

### Statistical analysis

All experiments were done with biological replicates and were performed at least two times. Statistical tests were performed using GraphPad Prism 8.0 (GraphPad Software, Inc., La Jolla, CA), and all data are represented as mean ± s.e.m. Group comparisons were assessed by Student’s *t*-test (two groups). All statistical tests were two-tailed, and *p* ≤ 0.05 was defined as significant.

## RESULTS

### Depletion of NCOR vs. SMRT triggers distinct transcriptome alterations

To elucidate the roles of NCOR and SMRT in gene regulation, we individually knocked down NCOR and SMRT in the mouse macrophage cell line RAW264.7 (hereafter for simplicity referred to as ‘RAW cells’). RNA-seq results demonstrated the significant reduction of NCOR and SMRT mRNA expression, while the mRNA expression of other complex subunits remained unaffected (**Supplementary Figure S1A**). Western blot analysis confirmed the efficient knockdown of NCOR and SMRT at the protein level (**Supplementary Figure S1B**). RNA-seq analysis revealed that the depletion of NCOR vs. SMRT, as compared to control cells, resulted in substantially different transcriptome alterations. Principal component analysis (PCA) identified a more prominent impact of SMRT depletion on the transcriptome, as compared to NCOR depletion (**Figure 1A**). We then conducted a differential gene expression analysis and classified the differentially expressed genes in six distinct clusters (**Figure 1B**). Cluster 1 comprises genes, such as *Trem2, Cd9*, and *Lgals3*, which were commonly upregulated upon depletion of both NCOR and SMRT. Cluster 2 consists of inflammatory and cancer-related genes, including *Cxcl2*, *Spp1* and *Ccl7,* upregulated upon SMRT depletion. Cluster 3 contains metabolic genes, such as *Cpt1a*, *Acads* and *Slc5a3,* upregulated upon NCOR depletion. The genes in cluster 5 that were downregulated upon SMRT depletion are relevant for oxidative phosphorylation (OXPHOS) (*Nduf*, *Uqcr* and *Sdh* family genes), but also for lipid metabolism and anti-inflammatory fatty acid production (*Srebf1* and *Srebf2,* encoding the TFs SREBP1/2 (56). Genes linked to tumor-associated macrophages (TAM) such as *Cd40* and *Cd274* (3,57) and genes related to viral infection (*Oas2*, *Tap1* and *Irf1*) were downregulated upon NCOR depletion. Lastly, it is noteworthy that NCOR and SMRT depletion exhibited even opposite effects on certain genes, such as *Il4i1* and *Pdcd1* (encoding for PD-L1, programmed cell death ligand), which are upregulated upon SMRT depletion but downregulated upon NCOR depletion, and conversely, *Abcg1* and *Cox5b* which are upregulated upon NCOR depletion but downregulated upon SMRT depletion (**Supplementary Figure S1C**). KEGG pathway analysis (**Figure 1C**, **Supplementary Figure S1D**) revealed that the genes that were commonly upregulated upon depletion of both NCOR and SMRT, as well as those specifically upregulated upon SMRT depletion or specifically downregulated upon NCOR depletion, are highly relevant to inflammation and tumorigenesis. Specific shNCOR-upregulated genes and shSMRT-downregulated genes exhibit functionality primarily associated with metabolic processes such as oxidative phosphorylation, fatty acid metabolism, and the citric acid cycle. Moreover, genes that depended on both NCOR and SMRT were mainly involved in cell cycle control. These results emphasize the collaborative yet distinct involvement of NCOR and SMRT in diverse biological processes within macrophages, rather than serving redundant functions.

**Figure 1.**
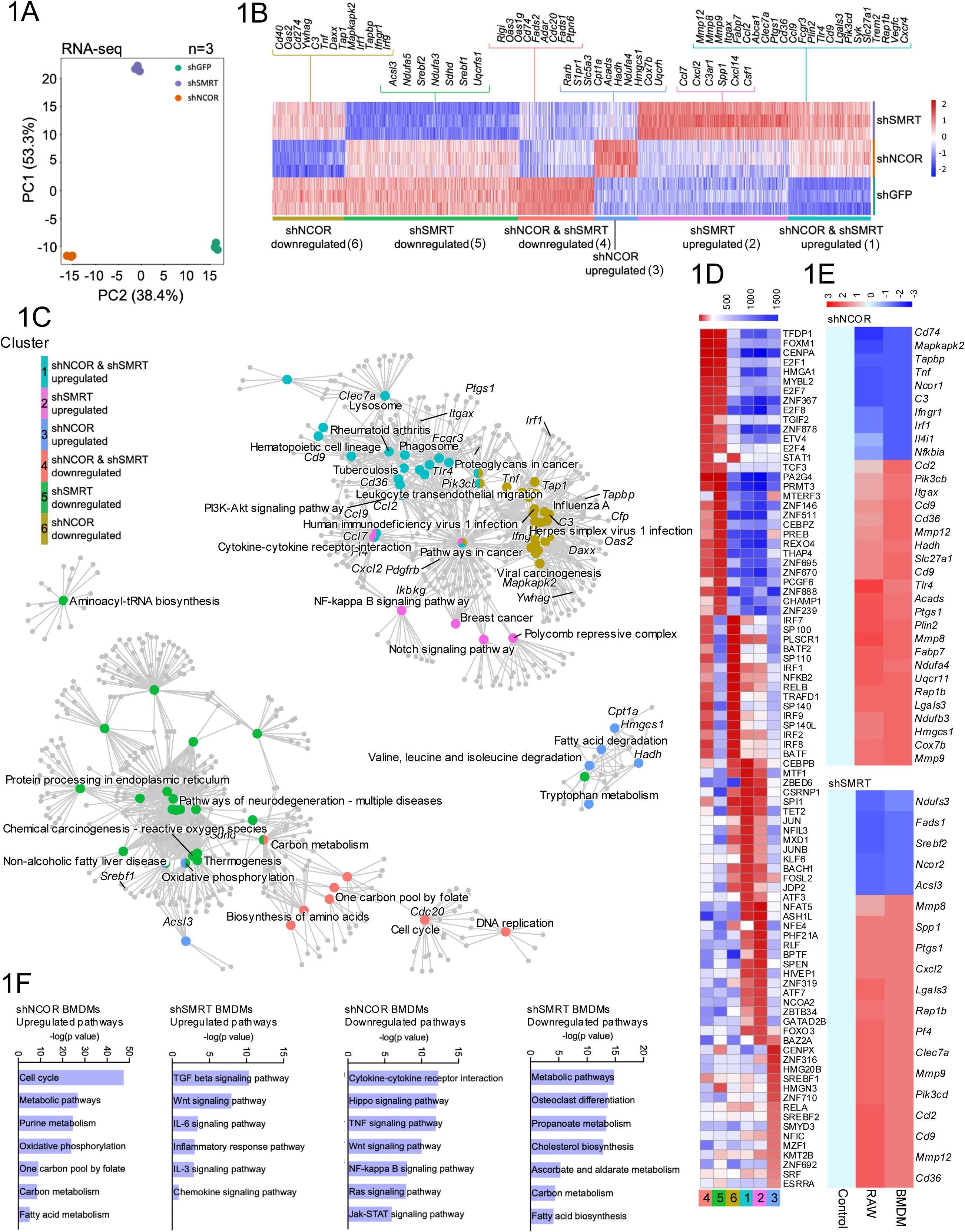
Depletion of NCOR vs. SMRT differentially alters gene expression (transcriptome). **(A)** Principal Component Analysis (PCA) illustrating the transcriptional outcomes in NCOR-vs. SMRT-depleted RAW cells (*n*=3). **(B)** Heatmap displaying differentially expressed genes in NCOR-vs. SMRT-depleted RAW cells, categorized in six clusters to show the different regulation patterns. Representative genes from each gene cluster are highlighted. Data significance for gene expression was determined using DESeq2. **(C)** Network of the top enriched KEGG pathways for each of the six gene clusters. **(D)** Heatmap showing the transcriptome-based TF activity analysis for each gene cluster. **(E)** Heatmaps illustrating selected genes in NCOR-depleted (top panel) and in SMRT-depleted (bottom panel) RAW cells and BMDMs. (F) Barplots showing enriched up- and down-regulated KEGG pathways in shNCOR BMDMs and shSMRT BMDMs.

To gain a more comprehensive understanding of the role of TFs in specifying NCOR vs. SMRT-mediated transcriptional regulation, we conducted a TF enrichment analysis using ChEA3 (45) within the aforementioned clusters. Our analysis unveiled the involvement of distinct TFs in each cluster (**Figure 1D**). In genes commonly upregulated by NCOR and SMRT depletion, or specifically upregulated by SMRT depletion, or specifically downregulated by NCOR depletion (clusters 1, 2 and 6 respectively), we identified lineage-determining TFs such as PU.1 (SPI1) and C/EBPB along with signal-responsive activating TFs such as JUN (an AP1 family member) (**Figure 1D**). However, cluster 6 is primarily regulated by inflammatory TFs such as IRFs and NFκB2. In cluster 3, comprising of genes specifically upregulated upon NCOR depletion, we observed lipid metabolism-regulating TFs such as SREBP1 and SREBP2 having increased activity (**Figure 1D**).

The gene regulation patterns and pathway alterations observed in RAW cells were also confirmed in primary mouse bone marrow-derived macrophages (BMDMs) treated with the same lentivirus shRNAs to knockdown SMRT vs. NCOR (**Figure 1E-F, Supplementary Figure S1F-G**). Additionally, the shRNA data were also confirmed in RAW cells treated with independent siRNA pools targeting different sequences than the shRNAs (**Supplementary Figure S1E, Supplementary Table S2)**.

All results together, in three independent knockdown approaches and two macrophage models, yield related differences in the gene expression networks altered upon depletion of NCOR (i.e. cells with remaining SMRT subcomplex) or depletion of SMRT (i.e. cells with remaining NCOR subcomplex), in comparison to control cells treated with shGFP/siCTRL (i.e. cells with intact NCOR-SMRT complex).

### Depletion of NCOR vs. SMRT differentially influences chromatin accessibility

To elucidate the role of the two corepressors in chromatin accessibility and remodeling, we conducted ATAC-seq (46,47) in NCOR- vs. SMRT-deficient RAW cells, as compared to control RAW cells. As in the transcriptome level **(Figure 1A)**, SMRT depletion has a more prominent effect than NCOR depletion in the accessibility of chromatin **(Figure 2A).** SMRT depletion resulted in more than 8000 differentially accessible peaks, with more than 5000 peaks upregulated. In contrast, NCOR depletion resulted in approximately 5000 differentially accessible peaks, with more than 2000 peaks upregulated. Chromatin accessibility was primarily upregulated in enhancer regions (**Figure 2B-C, Supplementary Figure S2A**). Integrating the ATAC-seq data with the corresponding RNA-seq data revealed a correlation between alterations in chromatin accessibility following the depletion of NCOR and SMRT and changes in gene transcription levels (**Figure 2D**, **2E**). We generated a heatmap of all differentially accessible regions (**Figure 2F**) to illustrate how corepressor depletion-dependent changes in chromatin accessibility mirror transcriptional changes (**Figure 1B**). Further analysis, integrating the ATAC-seq data with the NCOR and SMRT cistrome data, revealed that the corepressor binding sites aligned with accessible chromatin regions, suggesting the possible involvement of NCOR and SMRT in regulating chromatin accessibility at the occupied enhancer regions (**Supplementary Figure S2B-S2D**).

**Figure 2.**
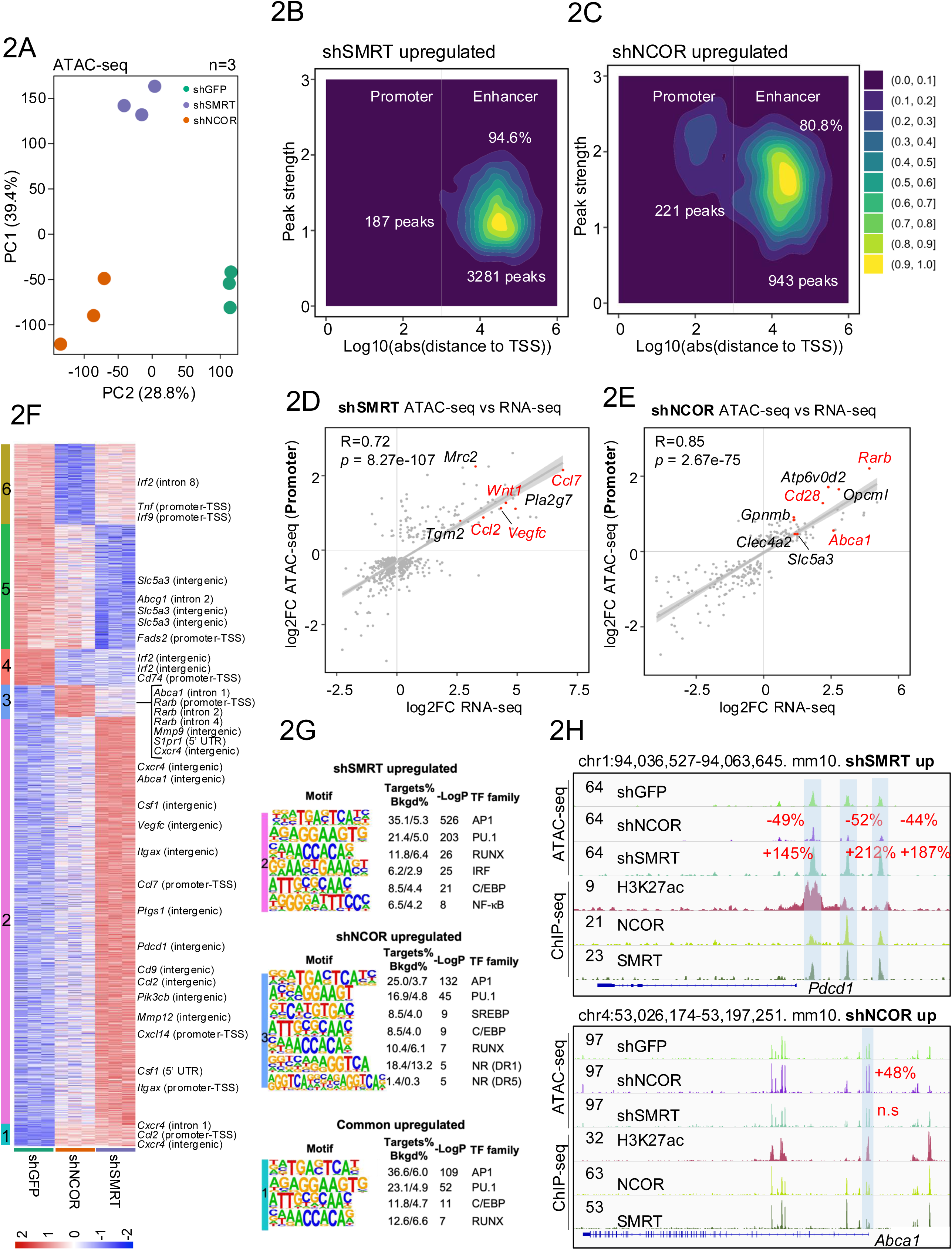
Depletion of NCOR vs. SMRT differentially alters chromatin accessibility (epigenome, candidate cis-regulatory elements). **(A)** PCA plot of ATAC-seq for NCOR-vs. SMRT- depleted cells (*n*=3). **(B-C)** Distribution of shSMRT- (**B**) shNCOR- (**C**) specific upregulated regions between enhancer and promoter regions. The distribution is presented along the distance from the transcription start site (TSS) of the annotated gene. (D-E) Scatter plots presenting the correlation of the RNA-seq expression and the promoter peaks from ATAC-seq data in SMRT- (**D**) and NCOR- (**E**) depleted cells. Data significance for gene expression was determined using DESeq2. **(F)** Heatmap of all differentially accessible genomic regions based on ATAC-seq data in NCOR- vs. SMRT- depleted cells, categorized in the same six clusters as in Figure 1B. Representative genomic regions from each cluster with differential accessibility are highlighted. (**G**) TF-binding site motif analysis of the SMRT-specific, NCOR- specific, and commonly repressed peaks. (**H**) IGV genome-browser tracks representing the NCOR, SMRT and H3K27ac ChIP-seq peaks in WT cells and the ATAC-seq changes in NCOR- vs. SMRT- depleted cells at the *Pdcd1* (top panel) and *Abca1* (bottom panel) loci. The statistically significantly changed peaks are highlighted with blue shadow.

TF motif analysis (**Figure 2G and Supplementary Figure S2E-S2G**) indicated that altered chromatin regions (i.e. up- or downregulated upon corepressor depletion) were highly enriched with AP1 and PU.1 motifs, consistent with these TF families being highly expressed key TFs in macrophages. Interestingly, in upregulated regions for both NCOR and SMRT the most enriched motif was AP1(**Figure S2G**), while in downregulated regions the most enriched motif was PU.1(**Supplementary Figure S2G**). Motifs for other TF families were also differentially enriched, though with higher *P*-values. For example, IRF motifs were enriched in shSMRT-upregulated regions and in shNCOR-downregulated regions, and SREBP and nuclear receptor (NR) motifs were enriched in shNCOR-upregulated regions. This suggests a possible role of above TFs in cooperating with the corepressor complex to modulate chromatin accessibility. Representative gene loci where accessibility is specifically upregulated upon SMRT depletion are *Pdcd1* (**Figure 2H**) and *Cxcr3* (**Supplementary Figure S2H**), those specifically altered upon NCOR depletion are *Abca1* (**Figure 2H**), and *Rarb* (**Supplementary Figure S2I**), and those commonly altered upon depletion of SMRT or NCOR are *Ptgs1* (**Supplementary Figure S2J**). We conclude that SMRT and NCOR modulate chromatin accessibility at enhancer regions of differentially and commonly regulated genes, thereby mechanistically linking corepressor-dependent chromatin remodeling to transcriptional regulation.

### Depletion of NCOR vs. SMRT differentially influences H3K27 acetylation at enhancers

Given that corepressor depletion-dependent chromatin remodeling primarily occured at enhancers, we expanded our epigenome analysis to H3K27ac, a key histone modification marking active enhancers linked to transcriptional activation (14,22). H3K27ac ChIP-seq revealed distinct alterations triggered by depletion of NCOR vs. SMRT (**Figure 3A**). SMRT depletion primarily upregulated enhancer acetylation linked to inflammatory genes, while NCOR depletion primarily upregulated enhancer acetylation linked to metabolic genes and downregulated enhancer acetylation linked to inflammatory genes (**Figure 3B**). Genome-wide H3K27ac peak heatmaps underscored the more pronounced impact of SMRT compared to NCOR (**Figure 3C, Supplementary Figure S3C**). Although shNCOR does not affect SMRT binding globally, our analysis identifies specific genomic regions where SMRT binding is dependent on NCOR. Notably, SMRT binding is significantly increased at 4,603 sites, such as Ccl2, and Tnf, upon NCOR depletion. Conversely, SMRT binding is significantly decreased at 5,477 sites, including Abca1 and Abcg1. Amongst the upregulated peaks in NCOR- vs. SMRT-depleted macrophages, only 11% were common (**Supplementary Figure S3A**). In regions upregulated upon NCOR depletion, 42% showed additional suppression of H3K27ac upon SMRT depletion, whereas in regions upregulated upon SMRT depletion, 30% showed increased suppression of H3K27ac upon NCOR depletion (**Supplementary Figure S3B**).

**Figure 3.**
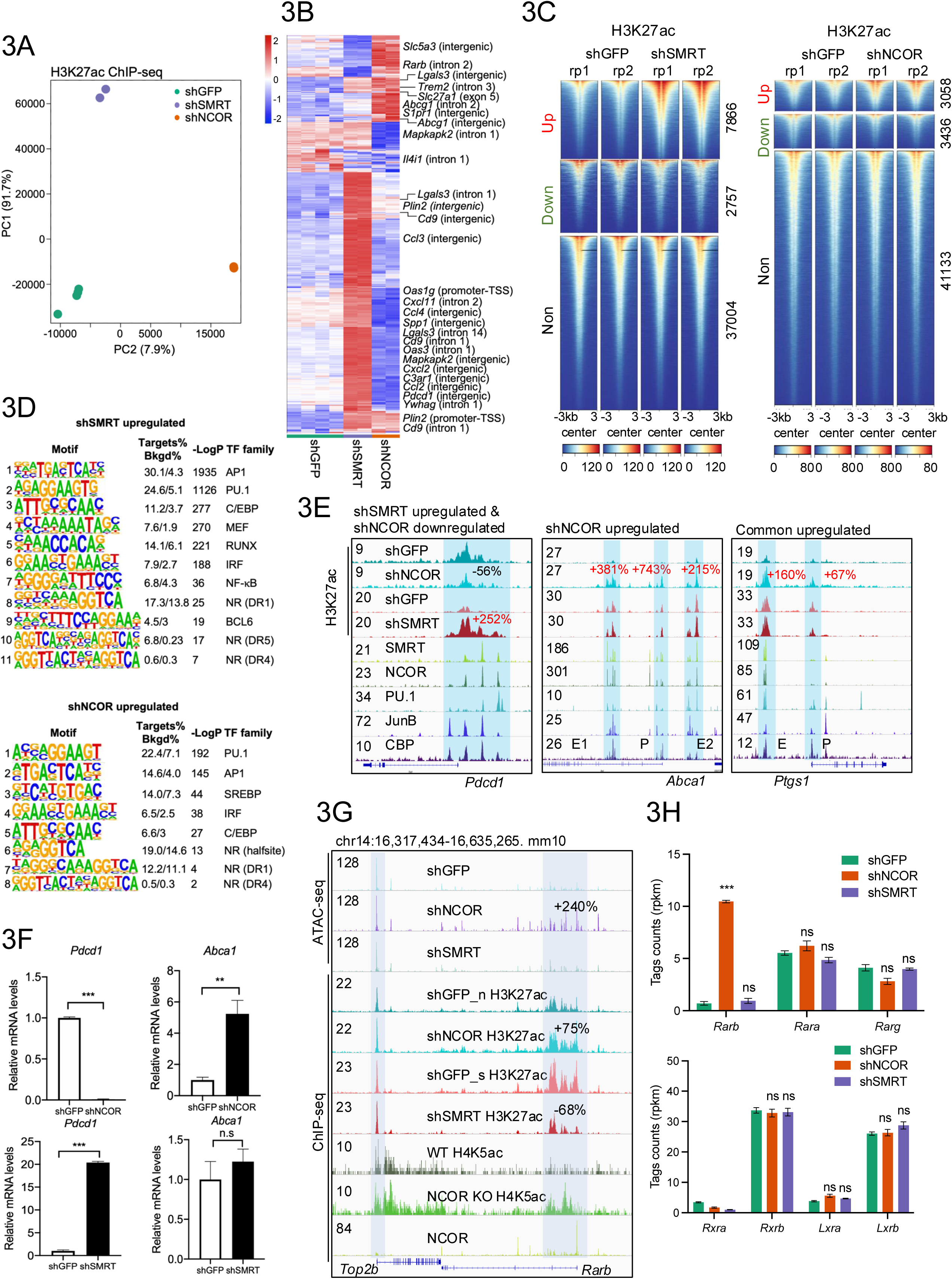
Depletion of NCOR vs. SMRT differentially alters H3K27ac (epigenome, enhancers). **(A)** PCA plot illustrating the H3K27ac ChIP-seq results for NCOR- vs. SMRT- depleted cells (*n*=2). Two independent ChIP-seq experiments were performed, and all results were merged to eliminate batch effect. **(B)** Heatmap displaying the z-normalized counts of the leading 2000 H3K27ac peaks driving PC1. Representative genomic regions are highlighted. **(C)** Heatmap of all H3K27ac peaks in shGFP, shSMRT and shNCOR macrophages. Up- and downregulated peaks are plotted independently (*n*=2). **(D)** Motif analysis of the upregulated H3K27ac peaks in NCOR- vs. SMRT- depleted cells, intersected with NCOR/SMRT peaks. **(E)** IGV genome browser tracks of H3K27ac ChIP-seq (basal condition) at *Pdcd1*, *Abca1* and *Ptgs1* loci. NCOR, SMRT, PU.1, JunB, and CBP ChIP-seq data are used to annotate enhancer regions. Up- and downregulated H3K27ac peaks are highlighted. **(F)** RT-qPCR analysis of *Pdcd1* and *Abca1* expression in NCOR- vs. SMRT- depleted cells (*n*=3). **(G)** IGV genome browser tracks of ATAC-seq and H3K27ac and H4K5ac ChIP-seq (basal condition) at the *Rarb* locus. Up- and downregulated peaks are highlighted with a blue shadow. **(H)** RNA-seq tag counts (-RPKM) of nuclear receptor gene expression in NCOR- vs. SMRT- depleted macrophages (*n*=3). Unpaired *t* test was used to determine data significance for gene expression in the qPCRs. All data are represented as mean ± SEM. Data significance for gene expression in RNA-seq was determined using DESeq2. \**P* < 0.05, \*\**P* < 0.01, \*\*\**P* < 0.001.

Focusing on regions marked by H3K27ac alterations upon depletion of NCOR or SMRT, we identified overlapping yet distinct TF motif enrichment patterns (**Figure 3D**). For example, AP1 was the most enriched motif in shSMRT-upregulated regions, followed by PU.1, C/EBP, MEF, RUNX, IRF, NF-κB and NR motifs, indicating a de-repression of these TFs upon SMRT depletion. In contrast, PU.1 was the most enriched motif in shNCOR-upregulated regions, followed by AP1, SREBP, IRF, C/EBP and NR motifs, while RUNX and NF-κB motifs were not enriched (**Figure 3D**). Representative H3K27ac changes were highlighted for corepressor-regulated genes including *Pdcd1*, *Abca1 (***Figure 3E)**, *Ccl3* and *Chst1* (**Supplementary Figure S3D, S3E**). H3K27ac changes were also observed in regions jointly upregulated by NCOR and SMRT depletion, such as the enhancer of *Ptgs1* (**Figure 3E**), consistent with the transcriptome and chromatin accessibility results. Interestingly, amongst the genes which were specifically upregulated by shNCOR but not shSMRT was *Rarb,* as supported by ATAC-seq, H3K27ac ChIP-seq, and RNA-seq data (**Figure 3G**, **3H**). *Rarb* encodes the retinoid acid receptor RAR beta, which heterodimerizes with RXRs and represents an established target TF for NCOR and SMRT (29). While well-studied in other cellular contexts, the function of RAR β in macrophages is currently unknown. Supporting the relevance of our RAW cell data, the activation-linked H4K5ac mark was increased, comparable to H3K27ac, at the *Rarb* locus in NCOR knockout BMDMs, as reported in an earlier study (**Figure 3G**) (38). As judged from our RNA-seq data, the expression of other RAR subtypes and additional relevant target nuclear receptors such as RXRs and LXRs was not affected upon NCOR or SMRT depletion (**Figure 3H**). In sum, these results demonstrate that depletion of NCOR or SMRT results in the differential alteration of H3K27ac levels at enhancers, consistent with the transcriptome changes.

### Cistrome analysis demonstrates genome-wide co-localization of NCOR and SMRT

Based on the existing literature, corepressors primarily affect transcriptional output through their interactions with CBP and H3K27ac. We present genome-scale data showing that NCOR, SMRT and CBP co-localize in macrophages and are associated with H3K27ac binding (**Supplementary Figure S4A**). Given the above identified differential roles of NCOR and SMRT in specifying the macrophage transcriptome and epigenome, we investigated whether differential chromatin binding patterns, might underlie these differences. This was not the case, as our analysis of the respective corepressor cistromes revealed a strong correlation between the binding sites of NCOR and SMRT (R=0.92, *p*<2.2e-16), indicating their significant genome-wide colocalization in RAW cells (**Figure 4A**). Approximately 75% of the corepressor binding sites were localized at enhancer regions, and 25% at promoter regions (**Figure 4B**). We then compared the cistromes of NCOR and SMRT with those of the myeloid lineage-determining TF PU.1 and the signaling-dependent TF JUNB (AP1 family), unveiling a robust colocalization with approximately 90% of their binding sites being shared (**Figure 4C, 4D**). We also found that NCOR and SMRT can predict the presence of super-enhancers (**Supplementary Figure S4B, S4C**), Additionally, we identified super-enhancer regions (**Supplementary Figure S4D-S4F**) labeled by NCOR and SMRT, either in common or specific regions.

**Figure 4.**
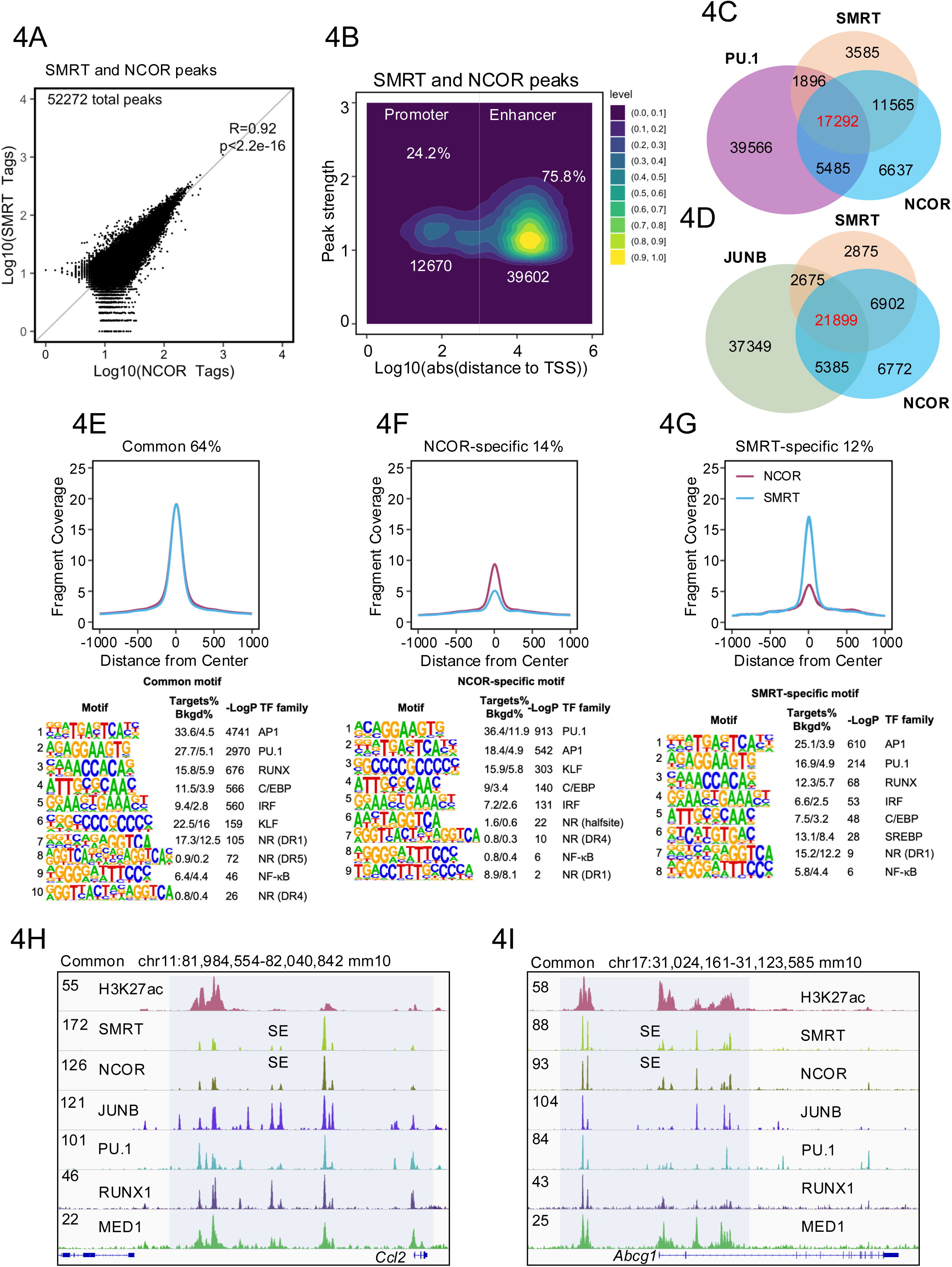
Genome-wide chromatin co-occupancy of NCOR and SMRT (cistrome). **(A)** Genome-wide correlation analysis: peaks from NCOR and SMRT ChIP-seq experiments were merged for Pearson correlation test. **(B)** NCOR and SMRT peak distribution between enhancer and promoter regions. The distribution is presented along the distance from the TSS of the annotated gene. **(C-D)** Venn diagrams displaying the binding overlap between NCOR, SMRT and the TFs PU.1 (**C**) and JunB (**D**) in macrophages. **(E-G)** Peak coverage plots of NCOR/SMRT common or specific peaks and the corresponding motif enrichment in macrophages. **(H-I)** IGV genome browser tracks of H3K27ac, SMRT, NCOR and other related regulatory proteins ChIP-seq at common marked genes *Ccl2* (**H**) and *Abcg1* (**I**) loci. The representative superenhancer (SE) and promoter regions of both *Ccl2* and *Abcg1* genes are highlighted with blue shadow.

Consistent with the genome-wide colocalization of NCOR and SMRT, DNA-binding motifs for PU.1 and AP1 were enriched in the corepressor-occupied regions (**Figure 4E**), suggesting the NCOR/SMRT-containing corepressor complex possibly to interact with these major macrophage TFs at chromatin. Notably, we observed distinct enrichments of 14% and 12% sites attributed specifically to NCOR or SMRT. While the functional relevance of these sites is uncertain, we note that the top-enriched motif was PU.1 for NCOR (**Figure 4F**) and AP1 for SMRT (**Figure 4G**). This possibly suggests that NCOR or SMRT, i.e. corepressor subcomplexes containing either NCOR or SMRT, exhibit in part distinct TF binding preferences, as compared to the ‘bona-fide corepressor complex’ containing both NCOR and SMRT (30). Representative for corepressor co-localization are the super-enhancer regions at the *Ccl2* and *Abcg1* loci (**Figure 4H, 4I**). In conclusion, our cistrome analysis reveals an extensive genome-wide colocalization of NCOR and SMRT in macrophages, making differential chromatin binding unlikely to account for the regulatory differences seen at the level of transcriptome and epigenome.

### Corepressor cistrome interdependence analysis identifies a superior role of SMRT

The chromatin co-occupancy of NCOR and SMRT does not inherently infer about their functional significance and interdependence. To further dissect their relationship at the cistrome level, we conducted ChIP-seq of NCOR and SMRT in RAW cells depleted for each corepressor. Remarkably, the cistrome data revealed a profound effect of SMRT depletion, as evidenced by the complete abolition of NCOR chromatin binding genome-wide, but not vice versa (**Figure 5A, 5B and Supplementary Figure S5A, S5B**). Visualization of specific gene loci supports above and suggests that NCOR does not substitute for the loss of SMRT to repress inflammatory genes *such as Ccl2 and Pdcd1* (**Supplementary Figure S5C**, **S5D**). While SMRT depletion also caused the dissociation of NCOR from metabolic gene loci such has *Abca1*, H3K27ac and gene activation was not affected (**Supplementary Figure S5E**).

**Figure 5.**
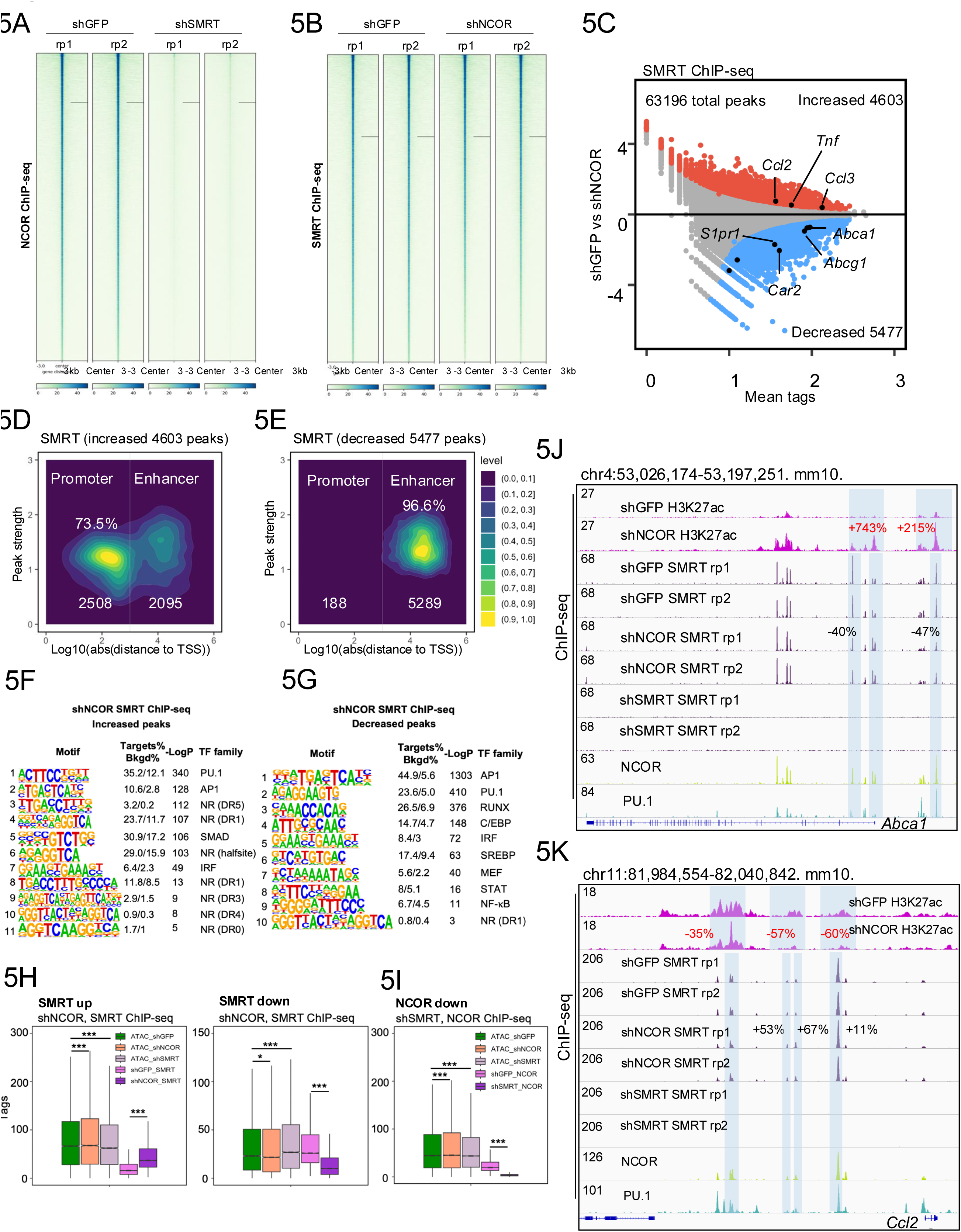
Reciprocal influence of NCOR and SMRT at the cistrome level. **(A-B)** Heatmap showing NCOR **(A)** and SMRT **(B)** occupancy in NCOR- vs. SMRT- depleted cells. The peak center in each plot represents the SMRT or NCOR peaks for each individual plot (*n*=2). **(C)** MA plot displaying SMRT peak changes in NCOR-depleted cells. Upregulated and downregulated peaks are highlighted (*n*=2). Key inflammatory and metabolic target genes are labeled. Significantly altered peaks were identified using DESeq2. **(D-E)** Distribution of up **(D)** and down **(E)** -regulated SMRT binding to promoters and enhancers in NCOR-depleted cells. The distribution is presented along the distance from the TSS of the annotated gene. **(F-G)** Motif analysis of up **(F)** and down **(G)** -regulated SMRT peaks in NCOR-depleted cells. **(H)** Boxplots comparing the distribution of the ATAC-seq tags between shNCOR/shSMRT and control and the SMRT ChIP-seq tags between shNCOR and control. The comparison is done for SMRT upregulated peaks (left panel) and for SMRT downregulated peaks (right panel). **(I)** Boxplots presenting the distribution of the ATAC-seq tags between shNCOR/shSMRT and control and the NCOR ChIP-seq tags between shSMRT and control. **(J-K)** IGV genome browser tracks showing SMRT ChIP-seq at *Abca1* and *Ccl2* loci. The corresponding H3K27ac ChIP-seq is used to illustrate the epigenome alterations (*n*=2). Wilcox test and DESeq2 were used to determine data significance for gene expression.

In strong support of the antibody specificity of the corepressor ChIP-seq, NCOR binding was completely abolished upon NCOR depletion, and SMRT binding was completely abolished upon SMRT depletion (**Supplementary Figure S5B).** Furthermore, our ChIP-seq analysis revealed a tighter association of GPS2 with SMRT rather than with NCOR. Upon SMRT depletion, GPS2 binding was nearly lost, whereas upon NCOR depletion GPS2 binding was reduced (**Supplementary Figure S5B**).

Remarkably, NCOR depletion resulted in a significant upregulation of 4603 SMRT peaks and a downregulation of 5477 SMRT peaks, indicating altered chromatin binding features of the remaining SMRT subcomplex lacking NCOR **(Figure 5C).** The upregulated SMRT peaks upon NCOR depletion were associated with inflammatory genes such as *Ccl2*, *Ccl3*, and *Tnf*, whereas the downregulated peaks were associated with metabolic genes such as *Abca1*, *Abcg1*, and *Car2* (**Figure 5C**). The changes in SMRT binding patterns correlated closely with changes in gene transcript levels (**Figure 1**). Subsequent analysis unveiled distinctive binding patterns for SMRT in the absence of NCOR. Specifically, upon NCOR depletion, the upregulated SMRT peaks were predominantly at promoters (**Figure 5D**), while downregulated SMRT peaks were predominantly at enhancers (**Figure 5E**). TF motif analysis revealed that peaks with increased SMRT binding upon NCOR depletion were highest enriched with motifs for PU.1, followed by AP1 and nuclear receptors (**Figure 5F).** In contrast, peaks with decreased SMRT binding upon NCOR depletion were highest enriched with motifs for AP1, followed by PU.1, RUNX, C/EBP and SREBP (**Figure 5G**). Further interestingly, there was a concurrent enhancement in chromatin openness, as defined by ATAC-seq, in regions where SMRT binding substantially increased upon NCOR depletion, but not in regions where SMRT binding was decreased (**Figure 5H**). This suggests that increased chromatin binding of SMRT (i.e. of a SMRT-containing subcomplex lacking NCOR) may influence chromatin accessibility. One the other hand, after SMRT is depleted NCOR binding is abolished genome-widely, but this is accompanied with a slight reduction in chromatin openness at a genome-wide level (**Figure 5I**). Our data also reveal reciprocal corepressor crosstalk at the epigenome and cistrome levels, exemplified by the metabolic *Abca1* gene locus where SMRT binding decreased upon NCOR depletion (**Figure 5J**), and the inflammatory *Ccl2* gene locus where SMRT binding increased upon NCOR depletion (**Figure 5K**).

### SMRT but not NCOR controls the nuclear localization of the corepressor complex

Given the requirement of SMRT but not of NCOR for the chromatin binding of the corepressor complex, we proceeded to further investigate the complex subunits at the subcellular level in RAW cells. Immunofluorescence (IF) analysis revealed that SMRT was present in nucleus of control lentivirus shGFP-treated cells as well as in NCOR-depleted cells treated with two different lentivirus shRNAs **(Figure 6A, S6A)**. In contrast, NCOR was present in nucleus only in control cells but accumulated in the cytoplasm of SMRT-depleted cells treated with two different lentivirus shRNAs **(Figure 6B, S6B)**. Nuclear exclusion of NCOR upon SMRT-depletion was similarly observed in BMDMs **(Figure 6C),** strongly supporting the conservation of this mechanisms in two mouse macrophage models.

**Figure 6.**
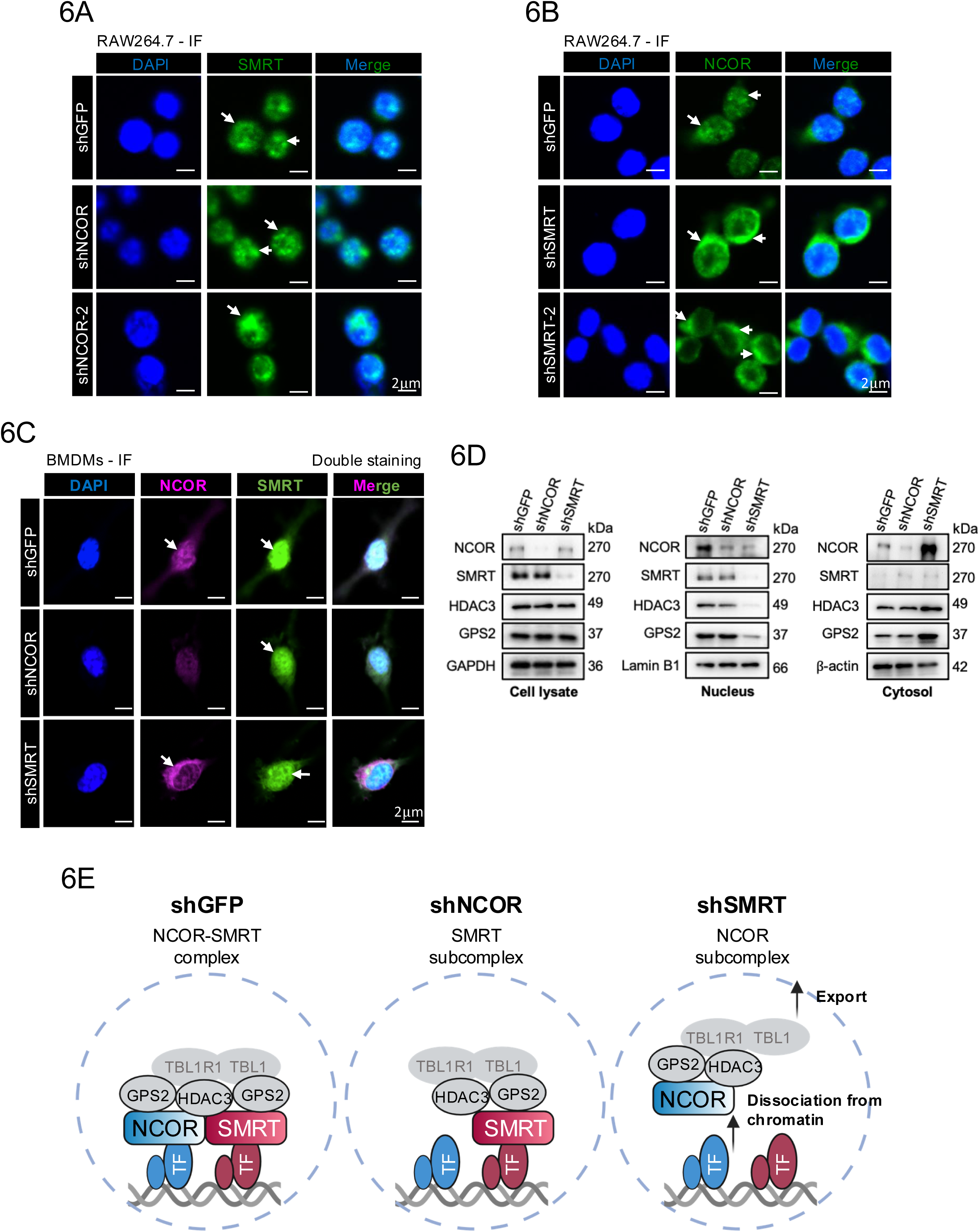
Influence of SMRT on NCOR subcellular localization. **(A)** Immunofluorescent (IF) staining showing the subcellular localization of SMRT in NCOR-depleted RAW cells; blue: DAPI, green: SMRT, scale bar: 2μm. **(B)** Immunofluorescent staining showing the subcellular localization of NCOR in SMRT-depleted RAW cells; blue: DAPI, green: NCOR, scale bar: 2μm. **(C)** Immunofluorescent double staining showing the subcellular localization of NCOR and SMRT in NCOR- and SMRT- depleted BMDMs; blue: DAPI, purple: NCOR, green: SMRT, scale bar: 2μm. **(D)** Subcellular fragmentation WB analysis of NCOR, SMRT, HDAC3 and GPS2 in NCOR- vs SMRT- depleted RAW cells. **(E)** Model of the compressor complex alterations upon NCOR vs. SMRT depletion in macrophages.

Complementary, we performed a subcellular fractionation analysis of the corepressor complex core subunits NCOR, SMRT, GPS2 and HDAC3 to follow their fate upon depletion of NCOR vs. SMRT in RAW cells (**Figure 6D)**. As judged from western blot analysis of whole cell extracts (**Figure 6D, left panel**), the protein levels of the remaining subunits were not altered upon depletion of either NCOR or SMRT. However, western blot analysis of nuclear extracts (**Figure 6D, middle panel**), revealed the loss of the remaining subunits upon depletion of SMRT, but not upon depletion of NCOR, fully consistent with the ChIP-seq results **(Figure 5A, Supplementary Figure S5B)**. Western blot analysis of cytosolic extracts **(Figure 6D, right panel)** revealed an accumulation of the remaining subunits (NCOR, HDAC3 and GPS2) in the cytosol, consistent with the immunofluorescence analysis (**Figure 6B**).

### Depletion of NCOR vs. SMRT differentially sensitizes macrophage activation by multiple signals

So far, our data suggested that depletion of NCOR or SMRT epigenetically remodels chromatin to transcriptionally reprogram unstimulated RAW cells, a state resembling M0 macrophages. To assess whether this epigenetic remodeling persists to influence signal-dependent macrophage activation, we stimulated corepressor-depleted RAW cells with lipopolysaccharide (LPS, a TLR4 activator triggering M1 activation (17)), IL4 (a STAT6 activator triggering alternative M2 activation (11,25) and GW3965 (a LXR agonist triggering metabolic activation and inhibiting inflammation (38,55,58–60)), followed by epigenome (H3K27ac CUT&Tag) and gene expression (RT-qPCR) analysis **(Figure 7 and Supplementary Figure 7**).

**Figure 7.**
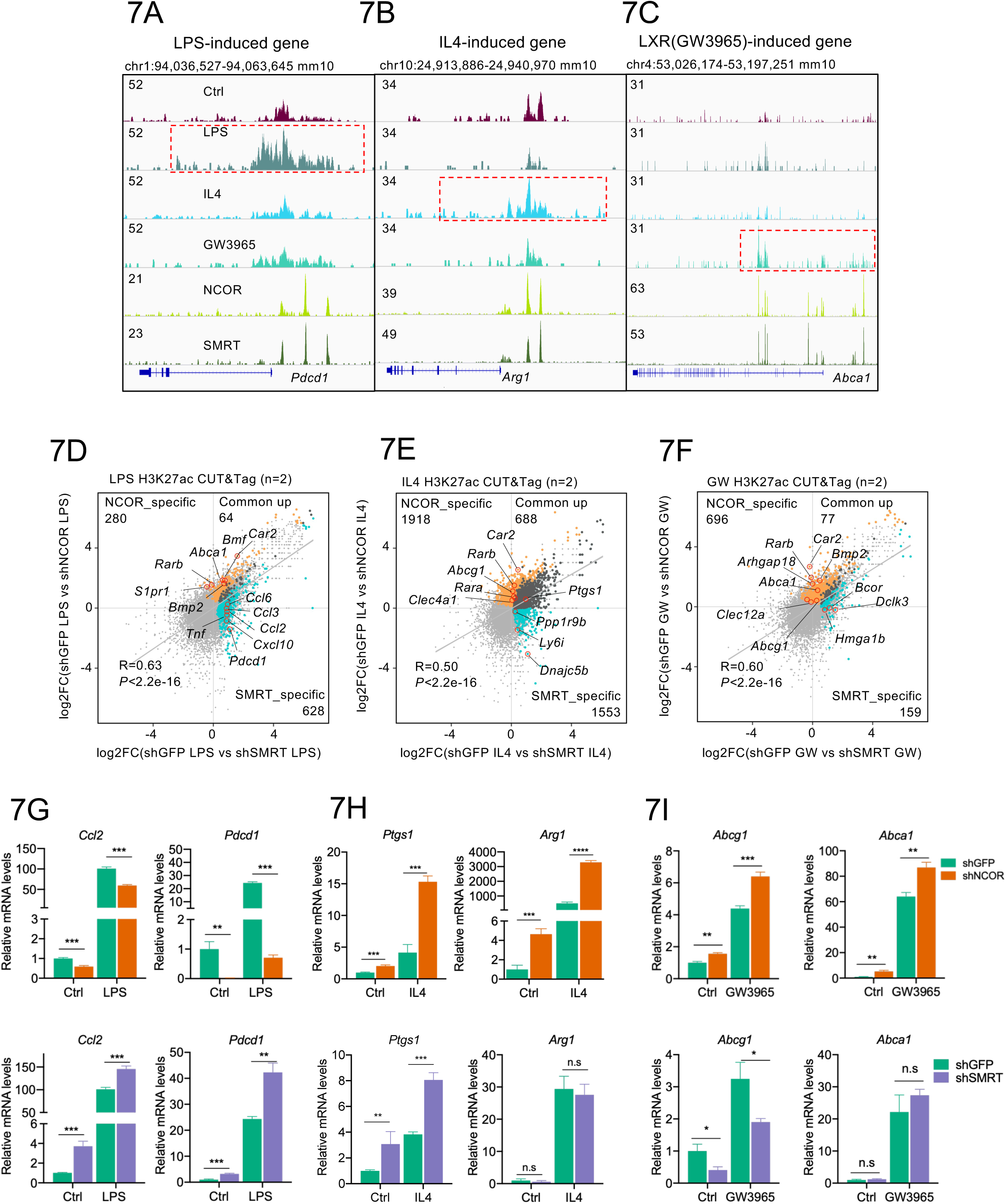
Depletion of NCOR vs. SMRT differentially reprograms macrophage activation in response to multiple signals. **(A-C)** IGV genome browser tracks displaying H3K27ac increase in *Pdcd1*, *Arg1*, and *Abca1* loci in response to LPS, IL4 or GW3965 treatment. Specific target epigenome regions are highlighted for each treatment. **(D-F)** Integration of CUT&Tag data for NCOR vs. SMRT depletion in LPS **(D)**, IL4 **(E)** or GW3965 **(F)** treated macrophages. The shNCOR-specific upregulated peaks are highlighted in orange and the shSMRT-specific upregulated peaks are highlighted in blue. **(G-I)** RT-qPCR analysis of related gene expression in LPS **(G)**, IL4 **(H)** and GW3965 **(I)** treatment in NCOR- vs. SMRT- depleted cells (*n*=3). Unpaired *t* test was used to determine data significance for gene expression. All data were represented as mean ± SEM. \**P* < 0.05, \*\**P* < 0.01, \*\*\**P* < 0.001, \*\*\*\**P* < 0.0001.

In control RAW cells expressing both corepressors, treatment with the corresponding stimuli induced H3K27ac-marked enhancer activation at representative response gene loci (*Pdcd1*, *Arg1*, *Abca1*) (**Figure 7A-7C**). The overall cellular state post-treatment reflected M1 (LPS), M2 (IL4), and metabolic activation (GW3965) (**Supplementary Figure S7A**-**S7C)**, with minimal overlap between the treatment responses (**Supplementary Figure S7D**). Interestingly, macrophage activation involves chromatin remodeling at enhancers and promoters to achieve gene regulation (**Supplementary Figure S7E**).

LPS/TLR4 activation of SMRT-depleted macrophages resulted in the increase of H3K27ac at regions linked to inflammatory genes, such as *Ccl2*, *Ccl3*, and *Pdcd1*, while NCOR depletion resulted in the increase of H3K27ac at regions linked to metabolic genes, such as *Abca1*, *Rarb*, *S1pr1*, and *Car2*. H3K27ac at the shNCOR-upregulated metabolic genes was further increased upon stimulation with IL4 or GW3965, whereas H3K27ac at the shSMRT-upregulated inflammatory genes was predominantly increased only upon LPS stimulation (**Figure 7D-7F**). Analysis of mRNA expression demonstrated that the depletion of NCOR or SMRT did not alter the levels of STAT6, JUN (AP1), or RELA/p65 (NF-kB) (**Supplementary Figure S7G**). Thus, NCOR and SMRT likely control the signal-induced transcriptional activity of these TFs at chromatin without affecting their expression. Finally, treating corepressor-depleted RAW cells with the corresponding stimuli revealed that mRNA expression changes of associated inflammatory, anti-inflammatory, and metabolic genes were concordant to H3K27ac alterations (**Figure 7G-7I**). Overall, these data suggest that the differential impact of NCOR vs. SMRT depletion on chromatin remodeling persists during signaling-dependent macrophage activation to specify gene expression programs characteristic for each remaining corepressor subcomplex (i.e the NCOR subcomplex upon SMRT depletion, the SMRT subcomplex upon NCOR depletion).

## DISCUSSION

Our study identifies critical non-redundant roles of SMRT and NCOR in differentially coordinating chromatin remodeling, histone acetylation, and transcription to regulate signal-dependent macrophage activation states linked to inflammation and metabolism. These differences are indicative of distinct functional subcomplexes and altered preferences to target TFs and to chromatin. We also uncover a hitherto unrecognized functional hierarchy with SMRT controlling the nuclear localization and the chromatin binding of the corepressor complex in macrophages.

For the interpretation of our comparative genome-wide and mechanistic SMRT vs. NCOR depletion analysis, we suggest that it is crucial to consider that SMRT and NCOR do not operate in isolation, i.e. without the remaining corepressor complex, to remodel chromatin and to regulate transcription. This consideration is based on the biochemical, structural and genome-wide evidence provided so far (for references, see Introduction), along with the lack of convincing experimental data suggesting that SMRT or NCOR also function complex-independently. Thus, the alterations triggered by depletion of SMRT or NCOR are likely reflecting the altered activities and target specificity of the respective remaining subcomplexes, i.e. the SMRT-deficient NCOR-containing subcomplex or the NCOR-deficient SMRT-containing subcomplex.

Our finding that SMRT depletion causes chromatin dissociation and nuclear export of NCOR, both in RAW cells and in BMDMs, is of particular interest. We propose a mechanistic model **(Figure 6)** in which macrophage SMRT serves as the nuclear anchor for NCOR and presumably for the remaining corepressor complex, as judged from our additional analysis of GPS2 and HDAC3. The relevance of the mechanism is supported by our genome-wide transcriptome and epigenome results, demonstrating that SMRT depletion has more pronounced effects than NCOR depletion on the number of differentially affected gene loci. Why NCOR dissociates from chromatin upon SMRT depletion is currently unclear and raises the question of how the chromatin binding of NCOR and SMRT is controlled in macrophages. Our data suggest that NCOR binding, in the absence of SMRT, is not simply determined by direct interactions with highly expressed macrophage TFs, including LXRs (38,61), BCL6 (36) and AP1 family members such as JUN and JUNB (62). Intracellular shuttling of NCOR and SMRT has been reported to be regulated by diverse signals, post-translational modifications and additional cofactors in different cell types (63–65) but remains to be explored in macrophages. The dominant role of SMRT in controlling NCOR, along with the associated core subunits HDAC3 and GPS2, is compatible with the intriguing possibility that one molecule SMRT and one molecule NCOR could be part of the same corepressor complex, as suggested by earlier biochemical and structural data (30). SMRT-depleted macrophages, along with related dendritic cell models (66,67), may be further explored as so far missing models of corepressor complex deficiency in the nucleus, complementary to HDAC3-directed PROTACs (68) and a double knockout model depleting both SMRT and NCOR, along with the complex, as recently described for hepatocytes (69).

Our study demonstrates that SMRT and NCOR, along with their respective subcomplexes, are pivotal coregulators that repress or activate substantially different inflammation-related gene expression networks in macrophages. This is especially evident upon TLR4 signaling triggering pro-inflammatory macrophage activation. SMRT functions anti-inflammatory by operating within a corepressor subcomplex along with GPS2 (17). However, NCOR functions pro-inflammatory by repressing anti-inflammatory LXR pathways (38) or by operating within an unusual coactivator complex along with HDAC3, PGC1β and p300/CBP (70). Based on our transcriptome analysis (i.e. down-regulated genes upon corepressor depletion), even SMRT exerts context-dependent coactivator-like activities, perhaps in cooperation with NCOR related to mechanisms most recently reported for hepatocytes (69). NCOR and SMRT subcomplexes exert significant effects on metabolic gene expression too, which often is coupled to anti-inflammatory pathways. NCOR but not SMRT seems critical for repressing fatty acid and cholesterol metabolism-related genes, many of which are regulated by LXRs (38). NCOR depletion additionally triggers activation of SREBP1, implicated in inflammation resolution by driving the synthesis of anti-inflammatory fatty acids (56). Additionally, we find that NCOR depletion upregulates OXPHOS-related genes, likely via activation of NRF1, consistent with previous studies that NCOR represses OXPHOS gene expression in muscle (71). In contrast, SMRT depletion downregulates electron transport chain genes and thereby impairs OXPHOS, consistent with previous findings using SMRT-mutated mouse models (40,41).

It remains to be seen whether the mechanisms underlying the non-redundant role of SMRT and NCOR also operate in other immune cell types such as dendritic cells, T cells or B cells, where related pathway alterations upon corepressor depletion have been reported (66,67,72,73). Notably, a comparative genomic and transcriptomic analysis in mouse dendritic cells demonstrated differential regulation of IL-6 and IL-10 by SMRT and NCOR (66), suggesting SMRT to act anti-inflammatory and NCOR pro-inflammatory just as in macrophages. Thus, the in part opposite regulatory roles of SMRT and NCOR subcomplexes in controlling inflammation-related pathways might be conserved between macrophages and dendritic cells, consistent with both myeloid cell types expressing overlapping sets of target TFs of the corepressor complex.

Mechanisms underlying the cell type-specific corepressor hierarchy remain to be further elucidated but may relate to the expression and composition of their respective target TFs in a given cell type, along with signaling-dependent posttranslational modifications of TFs and of the subunits of the corepressor complex, and possibly its subcomplexes. Evidence so far suggests that in macrophages highly expressed inflammatory TFs, such as the AP1 family members JUN and JUNB, and PU.1, recruit SMRT along with GPS2 to chromatin (17,25,37), indicative of a SMRT subcomplex. In contrast, in hepatocytes highly expressed metabolic nuclear receptors, such as the PPAR/RXR heterodimer, recruit NCOR along with GPS2, indicative of a NCOR subcomplex (74). Intriguingly, our comparative study suggests that the *in vivo* preferences of NCOR over SMRT subcomplexes for repressing nuclear receptors, such as LXRs PPAR and Rev-Erb, might be conserved in macrophages, consistent with earlier reports using NCOR-deficient mouse models (38,70,75,76).

Other mechanistic aspects of the role of NCOR and SMRT in determining corepressor sub-complex formation at chromatin remain elusive, such as details of their functional interplay with other core subunits. HDAC3 is the only enzymatic subunit of the complex due to its capability to deacetylate histones and other proteins, yet HDAC3 may also in part function independently of its catalytic activity (27,39,69,77,78) or even independently of the corepressor complex (39). While the precise role of HDAC3 in chromatin regulation and gene expression remains complex, evidence so far suggests that NCOR and HDAC3 share functions and cooperate in macrophages (27,78,79). GPS2 directly interacts with NCOR and SMRT but also with several TFs such as LXRs, PPARs, PU.1 and AP-1, as demonstrated by our previous studies (17,25,37,55,60,74). GPS2 has also been implicated in cytoplasmic signaling pathways (80–82), and it is not clear whether nuclear and cytoplasmic functions are mechanistically coupled. Notably, depletion of GPS2, unlike SMRT, has limited effects on chromatin accessibility, but mirrors SMRT depletion with respect to promoter-enhancer interactions, H3K27 acetylation and gene transcription (17,25). Finally, the coregulator exchange factors TBL1 and TBLR1 may also contribute to the regulation of corepressor subcomplex assembly, dynamics and chromatin binding (30,64,81,83).

As for the limitations of our study, we have not further dissected the mechanisms involved in transcriptional activation controlled by NCOR and SMRT, i.e. genes downregulated upon corepressor depletion. In case of SMRT, such mechanisms may be different from those recently reported for the NCOR-HDAC3-PGC1b ‘coactivator’ complex in TLR4-activated macrophages (70). They may perhaps relate to those most recently identified in hepatocytes and linked the regulation of chromatin accessibility and TF entry to chromatin (69). Further research is also needed to shed light on additional enigmatic functions of NCOR and SMRT, and their respective subcomplexes, possibly controlled by interactions with specific TFs, other coregulators and chromatin modifiers, or non-coding RNAs, along with diverse signal-dependent post-translational modifications. While our study utilized with RAW cells and BMDMs widely used mouse macrophage models *in vitro* and in part allows comparisons to previously published corepressor studies based on mouse models, the human conservation of key mechanisms and regulatory networks needs to be thoroughly investigated in future.

## DATA AVAILABILITY

RNA-seq, ChIP-seq, CUT&Tag, and ATAC-seq data generated in this study have been deposited at NCBI Gene Expression Omnibus (GEO) under the accession numbers GSE235408 and GSE291538. Public ChIP-seq/RNA-seq data generated by others were from GSE50944, GSE130383, GSE184884, and GSE247945 series.

## SUPPLEMENTARY DATA

Supplementary data are available online.

## ACKNOWLEDGEMENTS

We thank all members of our laboratories for sharing scientific materials and fruitful discussions. We acknowledge the BEA core facility (Karolinska Institutet) and Novogene for sequencing services. The computations were enabled by resources provided by the National Academic Infrastructure for Supercomputing in Sweden (NAISS).

## Authoŕs contributions

Z.H. and E.T. conceived the study. A.E., C.G., R.F. and Z.H. performed the wet lab experiments with contribution of Z.L. and O.G.; Z.H. and A.E. jointly performed the dry lab experiments and data analysis with contribution of all co-authors. A.E., Z.H. and E.T. wrote and edited the manuscript with input from all coauthors. Z.H. and E.T. acquired funding and jointly supervised the study.

