## Supplemental file for "Nonredundant roles of the HDAC3 corepressor complex subunits SMRT and NCOR in controlling inflammatory and metabolic macrophage pathways"

### Supplementary Information

**Efthymiadou et al.**

#### This file includes Supplementary Figures S1 to S7 and Supplementary Table S1 to S3.

### Figure S1

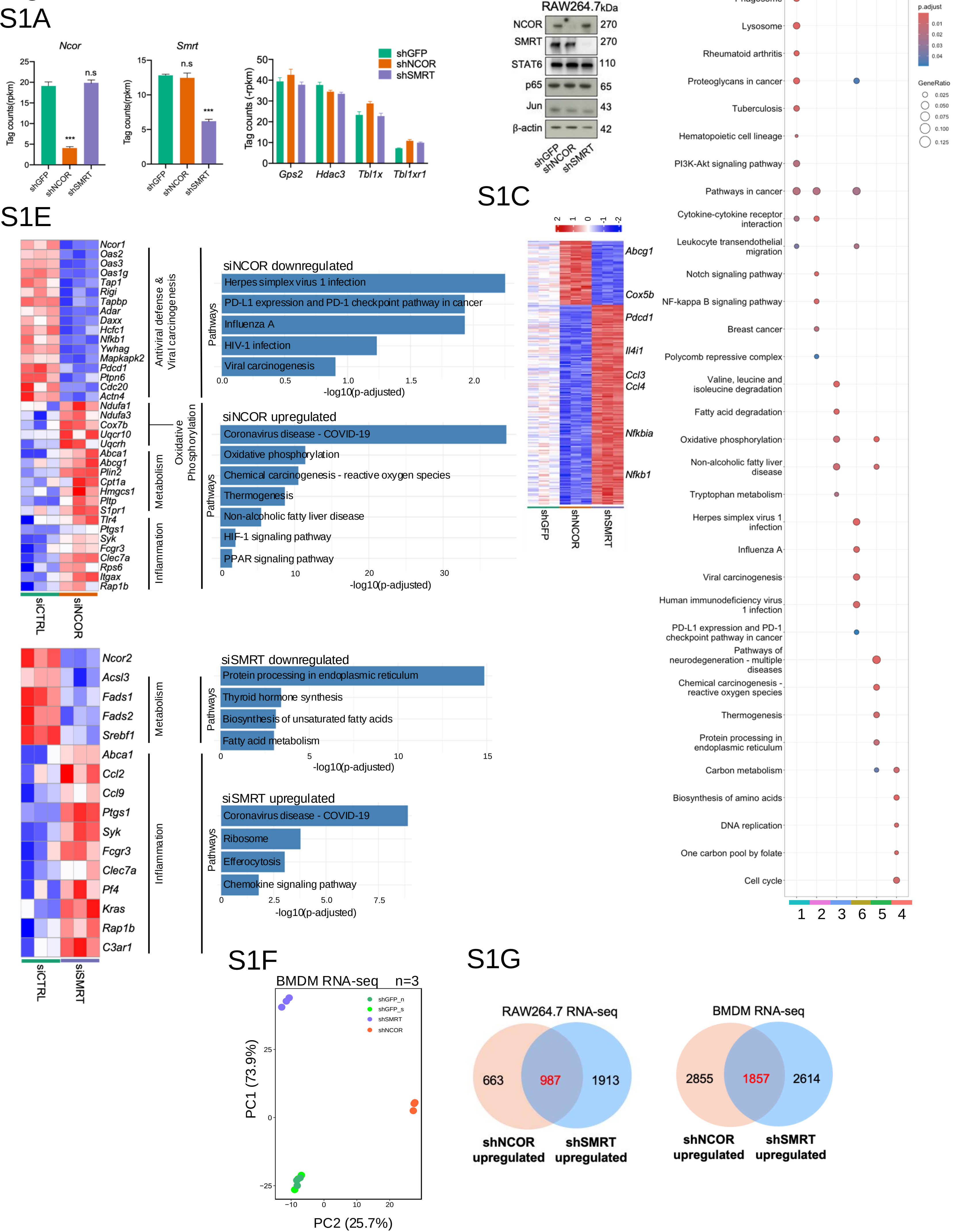
S1A
Ncor
Smrt
Tag counts(rpkm)
Tag counts (-rpkm)
shGFP
shNCOR
shSMRT
n.s
***
Gps2
Hdac3
Tbl1x
Tbl1xr1
S1B
RAW264.7
kDa
NCOR
SMRT
STAT6
p65
Jun
β-actin
270
270
110
65
43
42
S1C
Abcg1
Cox5b
Pdcd1
Il4i1
Ccl3
Ccl4
Nfkbia
Nfkb1
S1D
Phagosome
Lysosome
Rheumatoid arthritis
Proteoglycans in cancer
Tuberculosis
Hematopoietic cell lineage
PI3K-Akt signaling pathway
Pathways in cancer
Cytokine-cytokine receptor interaction
Leukocyte transendothelial migration
Notch signaling pathway
NF-kappa B signaling pathway
Breast cancer
Polycomb repressive complex
Valine, leucine and isoleucine degradation
Fatty acid degradation
Oxidative phosphorylation
Non-alcoholic fatty liver disease
Tryptophan metabolism
Herpes simplex virus 1 infection
Influenza A
Viral carcinogenesis
Human immunodeficiency virus 1 infection
PD-L1 expression and PD-1 checkpoint pathway in cancer
Pathways of neurodegeneration - multiple diseases
Chemical carcinogenesis - reactive oxygen species
Thermogenesis
Protein processing in endoplasmic reticulum
Carbon metabolism
Biosynthesis of amino acids
DNA replication
One carbon pool by folate
Cell cycle
1 2 3 6 5 4
p.adjust
GeneRatio
S1E
Antiviral defense & Viral carcinogenesis
Oxidative Phosphorylation
Metabolism
Inflammation
siCTRL
siNCOR
siSMRT
siNCOR downregulated
Herpes simplex virus 1 infection
PD-L1 expression and PD-1 checkpoint pathway in cancer
Influenza A
HIV-1 infection
Viral carcinogenesis
-log10(p-adjusted)
siNCOR upregulated
Coronavirus disease - COVID-19
Oxidative phosphorylation
Chemical carcinogenesis - reactive oxygen species
Thermogenesis
Non-alcoholic fatty liver disease
HIF-1 signaling pathway
PPAR signaling pathway
siSMRT downregulated
Protein processing in endoplasmic reticulum
Thyroid hormone synthesis
Biosynthesis of unsaturated fatty acids
Fatty acid metabolism
siSMRT upregulated
Coronavirus disease - COVID-19
Ribosome
Efferocytosis
Chemokine signaling pathway
Pathways
Ncor1 Oas2 Oas3 Oas1g Tap1 Rigi Tapbp Adar Daxx Hcfc1 Nfkb1 Ywhag Mapkapk2 Pdcd1 Ptpn6 Cdc20 Actn4 Ndufa1 Ndufa3 Cox7b Uqcr10 Uqcrh Abca1 Abcg1 Plin2 Cpt1a Hmgcs1 Pltp S1pr1 Tlr4 Ptgs1 Syk Fcgr3 Clec7a Rps6 Itgax Rap1b
Ncor2 Acsl3 Fads1 Fads2 Srebf1 Abca1 Ccl2 Ccl9 Ptgs1 Syk Fcgr3 Clec7a Pf4 Kras Rap1b C3ar1
S1F
BMDM RNA-seq n=3
shGFP_n
shGFP_s
shSMRT
shNCOR
PC1 (73.9%)
PC2 (25.7%)
S1G
RAW264.7 RNA-seq
663
987
1913
shNCOR upregulated
shSMRT upregulated
BMDM RNA-seq
2855
1857
2614

**Supplementary Figure S1.** Influence of NCOR vs. SMRT depletion on gene expression. **(A)** RNA-seq tag counts (-RPKM) of *Ncor* and *Smrt* gene expression (left panel) and of *Gps2*, *Hdac3*, *Tbl1x*, *Tbl1xr1* gene expression (right panel) in NCOR- vs. SMRT- depleted cells (*n*=3). **(B)** Western blot analysis of NCOR, SMRT and of the TFs p65, JUN and STAT6 in NCOR- vs. SMRT-depleted RAW cells. **(C)** Heatmap displaying differentially expressed genes in which NCOR and SMRT depletion have opposite effect. Representative genes from each gene cluster are highlighted. Data significance for gene expression was determined using DESeq2. **(D)** Selected enriched KEGG pathways for each of the six gene clusters. **(E)** Heatmaps of selected differentially expressed genes in siNCOR and siSMRT RAW cells (*n*=3) (left panel) and barplots of their respective enriched KEGG pathways (right panel). (**F**) PCA plot illustrating the transcriptional outcomes in NCOR- vs. SMRT-depleted BMDMs (*n*=3). (**G**) Venn diagrams showing the common genes that are upregulated in shNCOR and shSMRT in BMDMs (right panel) and in RAW cells (left panel). Data significance for gene expression was determined using DESeq2.

Figure S2

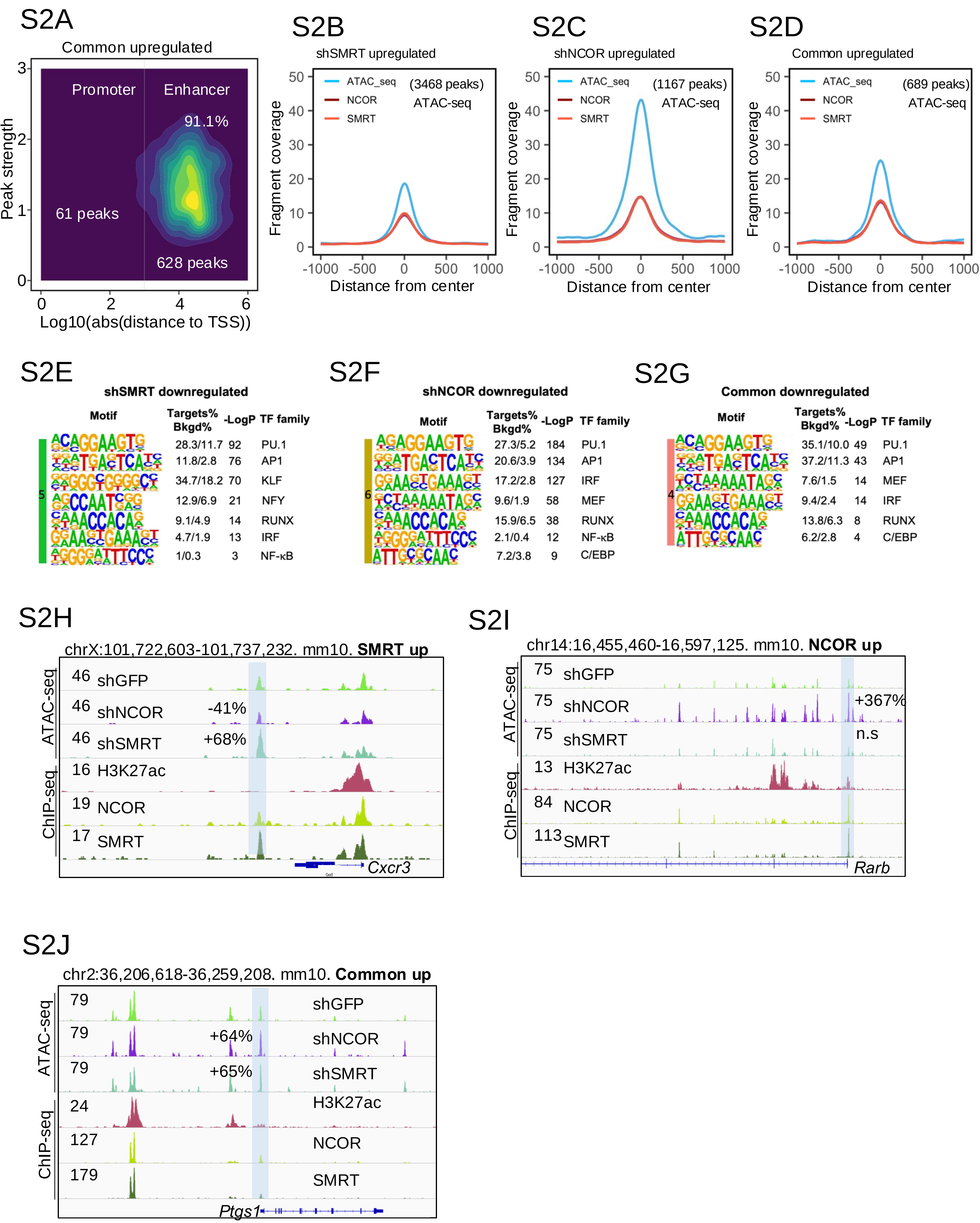
S2A
Common upregulated
Promoter
Enhancer
91.1%
61 peaks
628 peaks
Peak strength
Log10(abs(distance to TSS))
S2B
shSMRT upregulated
(3468 peaks)
ATAC-seq
S2C
shNCOR upregulated
(1167 peaks)
ATAC-seq
S2D
Common upregulated
(689 peaks)
ATAC-seq
ATAC_seq
NCOR
SMRT
Fragment coverage
Distance from center
S2E
shSMRT downregulated
Motif
Targets% Bkgd%
-LogP
TF family
28.3/11.7 92 PU.1
11.8/2.8 76 AP1
34.7/18.2 70 KLF
12.9/6.9 21 NFY
9.1/4.9 14 RUNX
4.7/1.9 13 IRF
1/0.3 3 NF-κB
S2F
shNCOR downregulated
27.3/5.2 184 PU.1
20.6/3.9 134 AP1
17.2/2.8 127 IRF
9.6/1.9 58 MEF
15.9/6.5 38 RUNX
2.1/0.4 12 NF-κB
7.2/3.8 9 C/EBP
S2G
Common downregulated
35.1/10.0 49 PU.1
37.2/11.3 43 AP1
7.6/1.5 14 MEF
9.4/2.4 14 IRF
13.8/6.3 8 RUNX
6.2/2.8 4 C/EBP
S2H
chrX:101,722,603-101,737,232. mm10. SMRT up
ATAC-seq
ChIP-seq
46 shGFP
46 shNCOR -41%
46 shSMRT +68%
16 H3K27ac
19 NCOR
17 SMRT
Cxcr3
S2I
chr14:16,455,460-16,597,125. mm10. NCOR up
75 shGFP
75 shNCOR +367%
75 shSMRT n.s
13 H3K27ac
84 NCOR
113 SMRT
Rarb
S2J
chr2:36,206,618-36,259,208. mm10. Common up
79 shGFP
79 shNCOR +64%
79 shSMRT +65%
24 H3K27ac
127 NCOR
179 SMRT
Ptgs1

**Supplementary Figure S2.** Influence of NCOR vs. SMRT depletion on chromatin accessibility.

**(A)** Distribution of commonly upregulated regions by SMRT and NCOR KD. The distribution is presented along the distance from the TSS of the annotated gene. **(B-D)** Peak coverages showing the overlap between NCOR and SMRT -binding regions and open chromosomal regions (ATAC-seq). SMRT-specific (**B**), NCOR-specific (**C**) and common (**D**) binding regions were plotted against ATAC-seq peaks. **(E-G)** Motif analysis of the shSMRT-specific (**E**), shNCOR-specific (**F**), and common (**G**) downregulated peaks. (**H-J**) IGV genome-browser tracks displaying the NCOR, SMRT, and H3K27ac ChIP-seq peaks in WT cells and the ATAC-seq changes in NCOR- vs. SMRT- depleted cells at *Cxcr3* (**H**), *Rarb* (**I**), and *Ptgs1* (**J**) loci. The statistically significantly changed peaks are highlighted with blue shadows. Data significance for peaks was determined using DESeq2.

### Figure S3

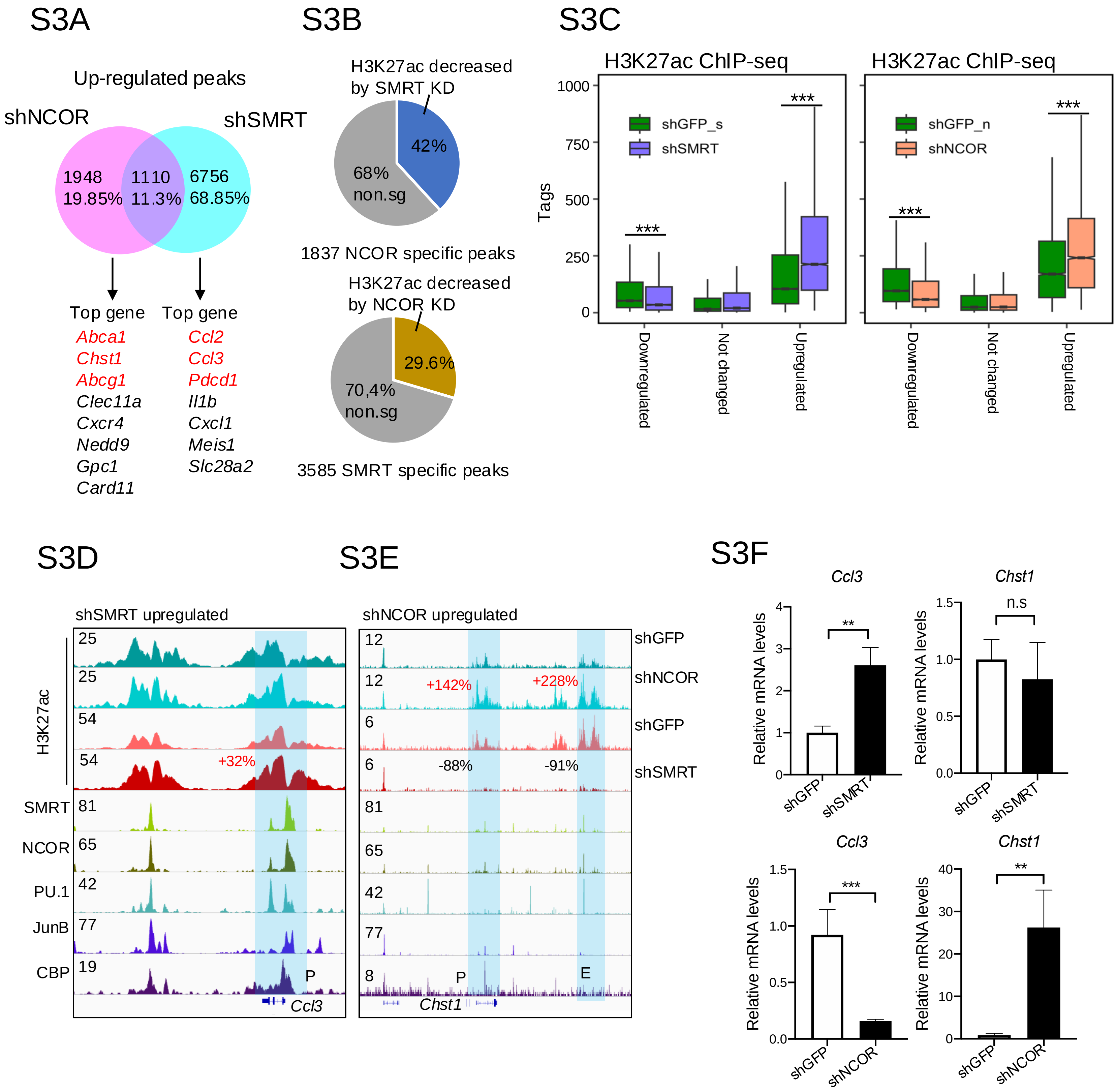

S3A
Up-regulated peaks
shNCOR
shSMRT
1948
19.85%
1110
11.3%
6756
68.85%
Top gene
Abca1
Chst1
Abcg1
Clec11a
Cxcr4
Nedd9
Gpc1
Card11
Top gene
Ccl2
Ccl3
Pdcd1
Il1b
Cxcl1
Meis1
Slc28a2
S3B
H3K27ac decreased by SMRT KD
42%
68% non.sg
1837 NCOR specific peaks
H3K27ac decreased by NCOR KD
29.6%
70,4% non.sg
3585 SMRT specific peaks
S3C
H3K27ac ChIP-seq
shGFP_s
shSMRT
Tags
shGFP_n
shNCOR
Downregulated
Not changed
Upregulated
S3D
shSMRT upregulated
H3K27ac
SMRT
NCOR
PU.1
JunB
CBP
+32%
Ccl3
S3E
shNCOR upregulated
+142%
+228%
-88%
-91%
Chst1
shGFP
shNCOR
shGFP
shSMRT
S3F
Ccl3
Chst1
Relative mRNA levels
shGFP
shSMRT
shNCOR
n.s

**Supplementary Figure S3.** Influence of NCOR vs. SMRT depletion on H3K27ac enhancer modification.

**(A)** Venn diagram depicting the overlap of the upregulated H3K27ac peaks in NCOR- vs. SM RT- depleted cells. Representative genes are highlighted for each cluster. **(B)** Pie charts illustrating the proportion of functional roles controlled by NCOR or SMRT in specific H3K27ac activation regions in corepressor-depleted cells. (**C**) Boxplots comparing the statistical difference of the H3K27ac ChIP-seq tags between shSMRT and control in shSMRT-downregulated, -upregulated and not changed peaks (left panel) and between shNCOR and control in shNCOR-downregulated, -upregulated and not changed peaks (right panel). **(D-E)** IGV genome browser tracks for H3K27ac ChIP-seq (basal condition) at the *Ccl3* (SMRT-repressed) and *Chst1* (NCOR-repressed) loci. Upregulated and downregulated H3K27ac peaks are highlighted. **(F)** RT-qPCR analysis of *Ccl3* and *Chst1* expression in NCOR- vs. SMRT-depleted cells ($n$=3). Unpaired $t$-test was used to determine data significance for gene expression. Wilcox test was used to determine data significance for the sequencing results. All data are represented as mean ± SEM. *$P < 0.05$, **$P < 0.01$, ***$P < 0.001$.

Figure S4

S4A

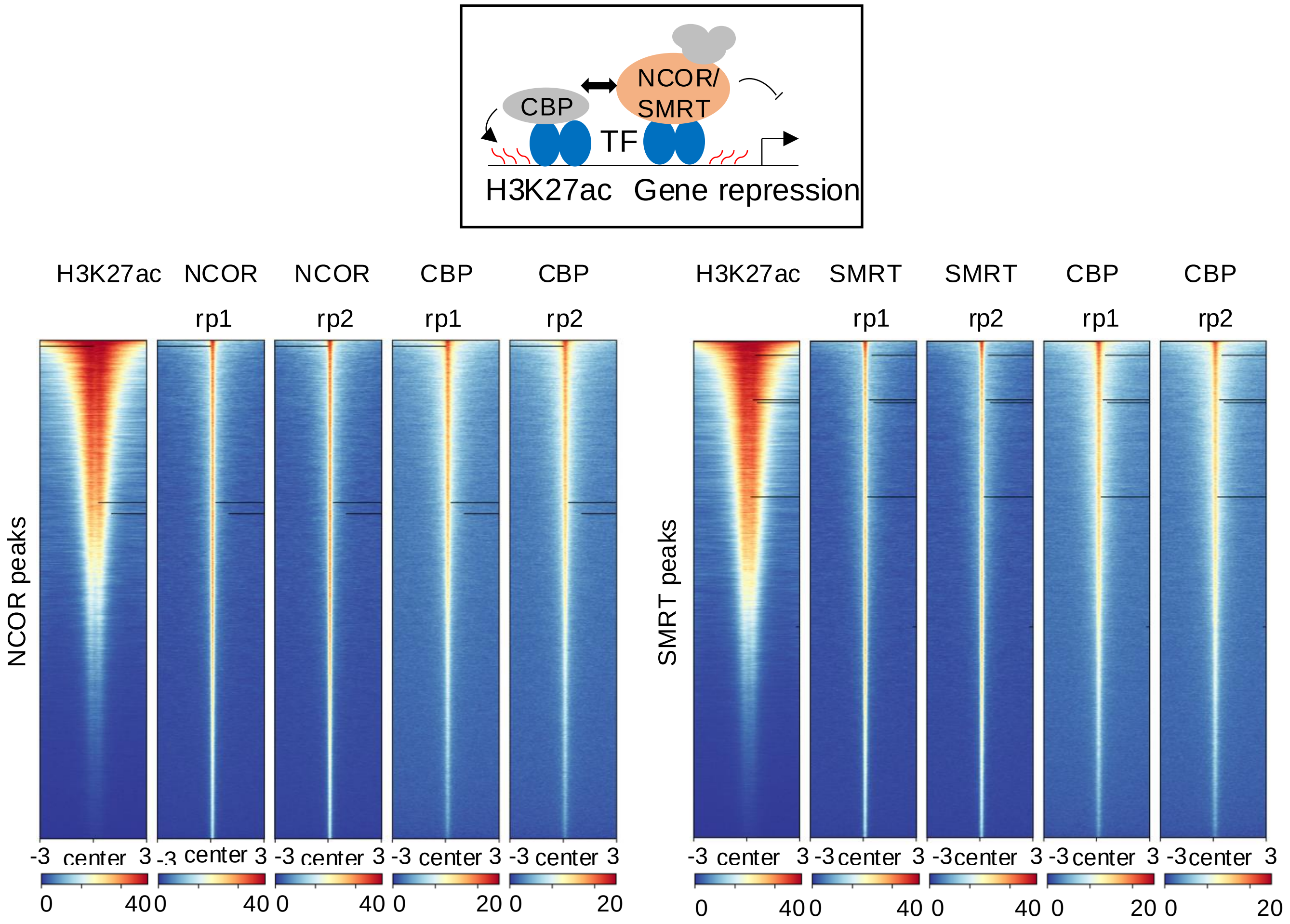


S4B

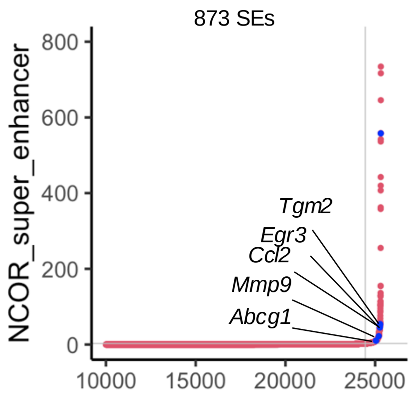


S4C

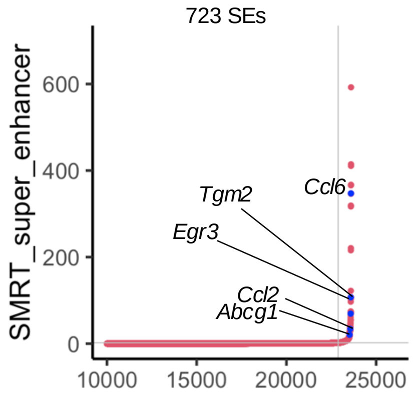


S4D

Common SE

chr2:158,098,837-158,179,438. mm10

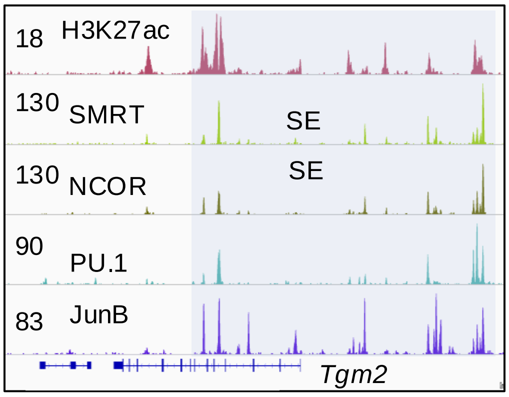


S4E

NCOR SE

chr1:135,605,464-135,761,857. mm10

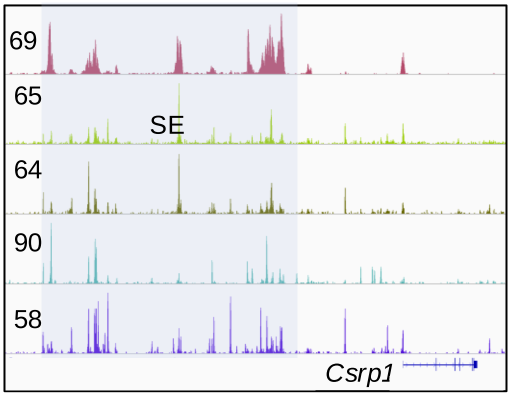


S4F

SMRT SE

chr1:191,165,447-191,263,209. mm10

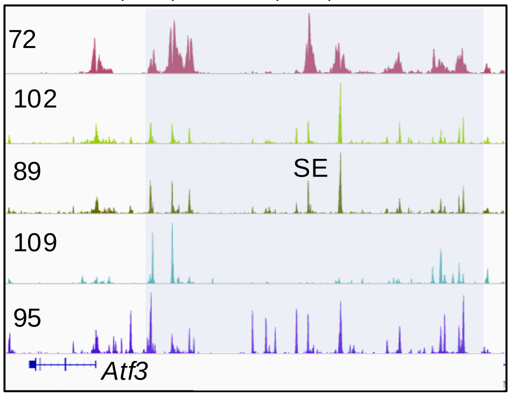

**Supplementary Figure S4.** Analysis of H3K27ac super-enhancers.
**(A)** Illustration of the antagonism between the NCOR/SMRT corepressors and the fundamental CBP/P300 coactivators in regulating H3K27ac activation during macrophage activation. The upper panel illustrates the regulatory model. The lower panel shows the heatmaps of H3K27ac and NCOR, SMRT and CBP ChIP-seq global binding patterns. **(B-C)** Ranking of enhancer regions associated with NCOR (**B**) and SMRT (**C**) based on the abundance of their respective ChIP-seq signals. Selected super-enhancers (SE) are highlighted. **(D-F)** IGV genome browser tracks of selected gene regions demonstrating the presence of super-enhancer regions that are common (**D**), NCOR-specific (**E**), SMRT-specific (**F**).

### Figure S5

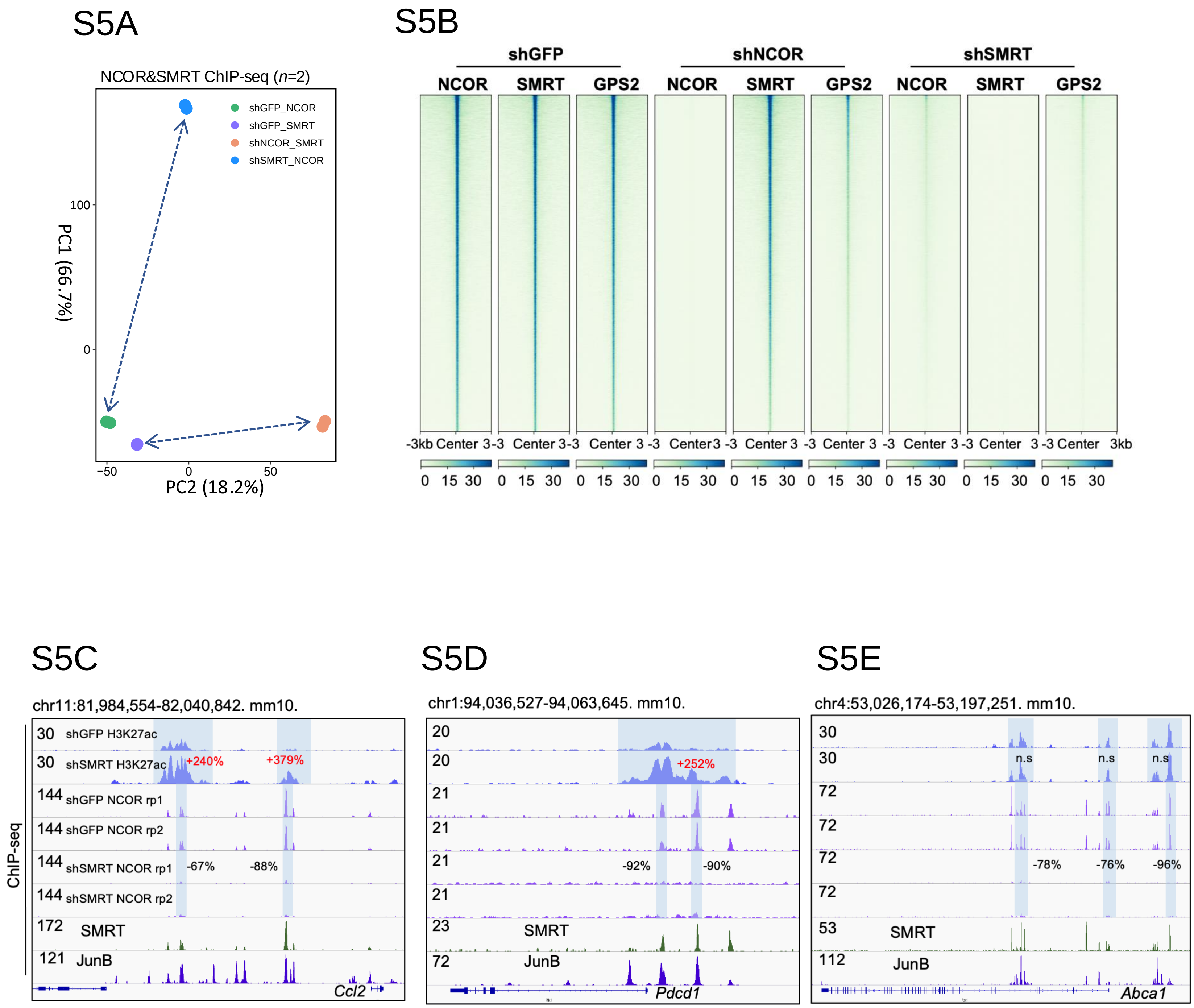
S5A
NCOR&SMRT ChIP-seq (n=2)
shGFP_NCOR
shGFP_SMRT
shNCOR_SMRT
shSMRT_NCOR
PC1 (66.7%)
PC2 (18.2%)
100
0
−50
0
50
S5B
shGFP
shNCOR
shSMRT
NCOR
SMRT
GPS2
-3kb Center 3kb
0 15 30
S5C
chr11:81,984,554-82,040,842. mm10.
ChIP-seq
30 shGFP H3K27ac
30 shSMRT H3K27ac
+240%
+379%
144 shGFP NCOR rp1
144 shGFP NCOR rp2
144 shSMRT NCOR rp1
-67%
-88%
144 shSMRT NCOR rp2
172 SMRT
121 JunB
Ccl2
S5D
chr1:94,036,527-94,063,645. mm10.
20
20
+252%
21
21
21
-92%
-90%
21
23 SMRT
72 JunB
Pdcd1
S5E
chr4:53,026,174-53,197,251. mm10.
30
30
n.s
n.s
n.s
72
72
72
-78%
-76%
-96%
72
53 SMRT
112 JunB
Abca1

**Supplementary Figure S5.** Analysis of NCOR vs. SMRT cistromes (chromatin binding). **(A)** PCA plot of NCOR and SMRT cistrome binding (via ChIP-seq) in NCOR- vs. SMRT-depleted cells (*n*=2). **(B)** Heatmaps of ChIP-seq peaks revealing the cistrome dependence of the three TF-binding corepressor complex subunits NCOR, SMRT and GPS2 in control (shGFP), NCOR-depleted cells and SMRT-depleted cells (*n*=2). **(C-E)** IGV genome browser tracks showing NCOR binding on *Ccl2* (**C**), *Pdcd1* (**D**), and *Abca1* (**E**) gene loci in SMRT-depleted cells, annotated with corresponding H3K27ac results.

### Figure S6

## S6A

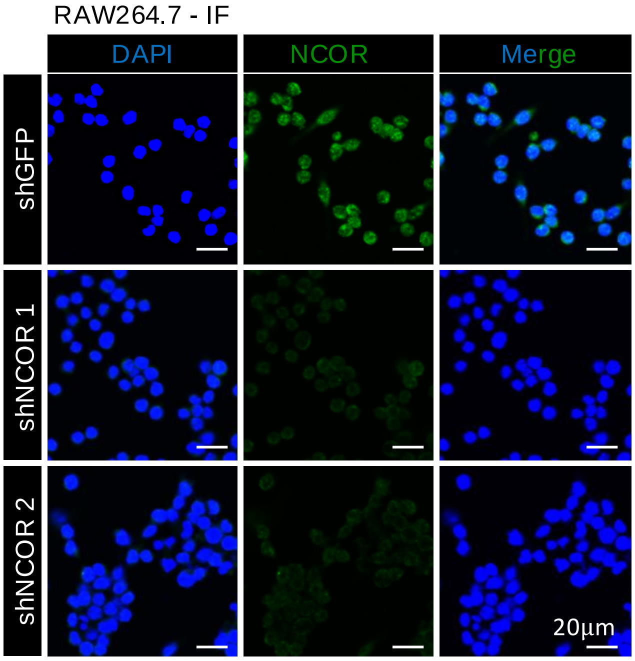


## S6B

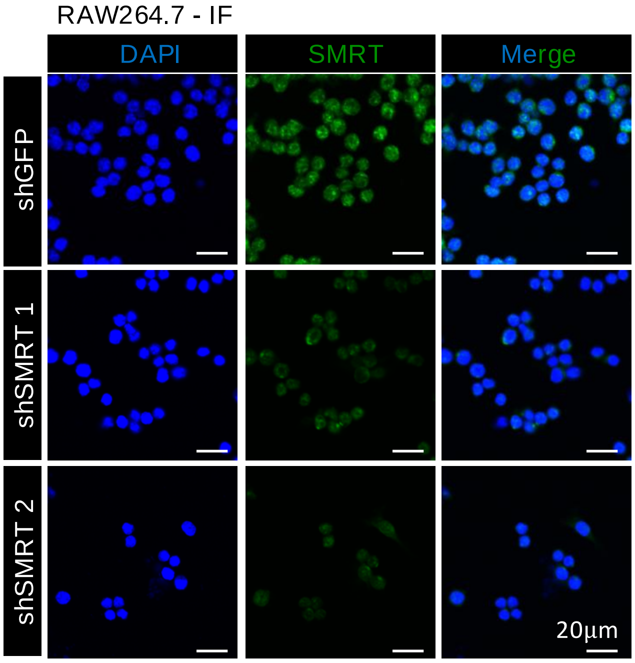

**Supplementary Figure S6.** Influence of SMRT on NCOR subcellular localization. **(A)** Immunofluorescent stainings showing the efficient removal of NCOR in NCOR-depleted RAW cells; blue: DAPI, green: NCOR, scale bar: 20μm. **(B)** Immunofluorescent staining showing the efficient removal of SMRT in SMRT-depleted RAW cells; blue: DAPI, green: SMRT, scale bar: 20μm.

### Figure S7

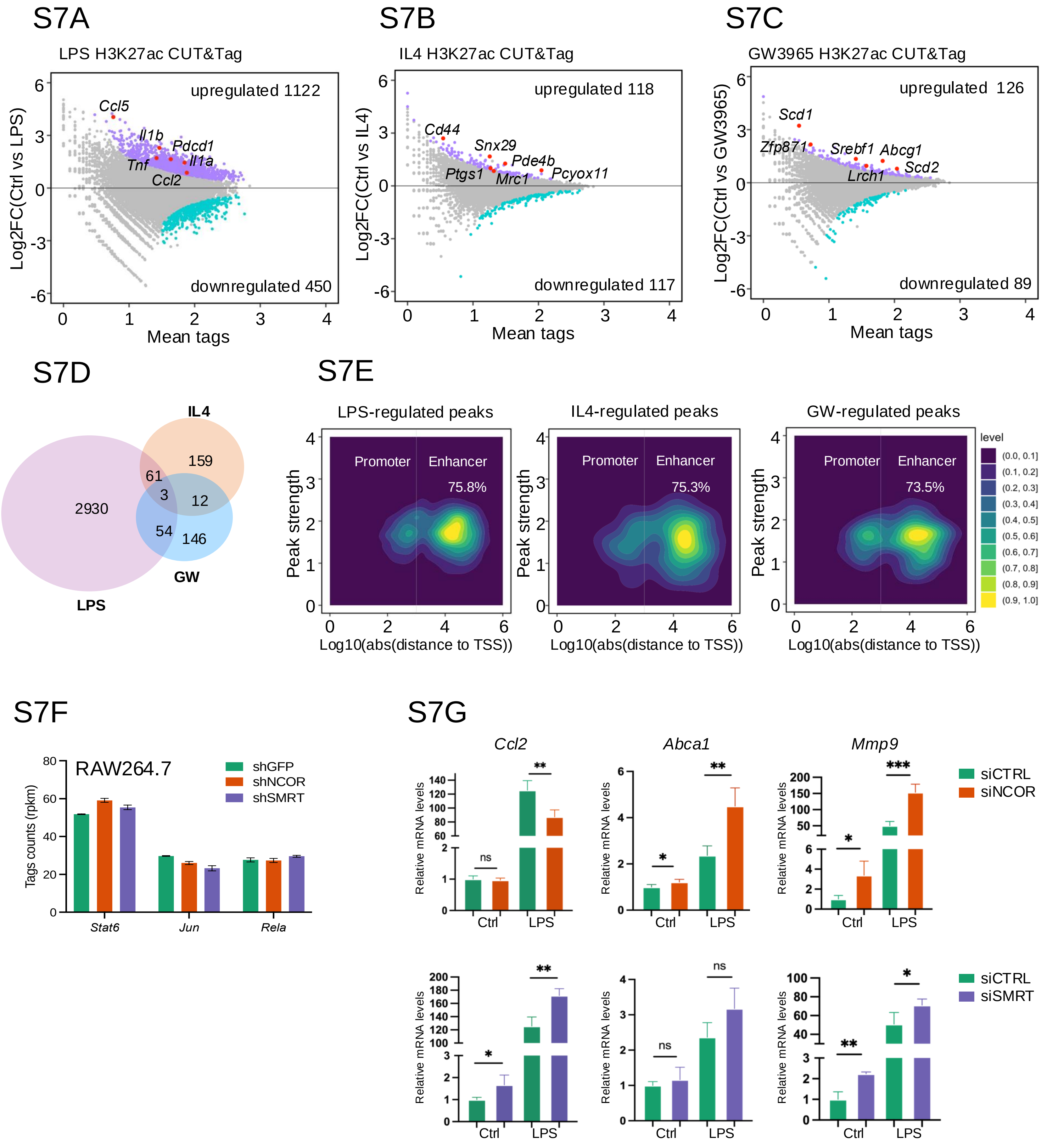
S7A
LPS H3K27ac CUT&Tag
upregulated 1122
downregulated 450
Ccl5
Il1b
Pdcd1
Il1a
Tnf
Ccl2
Log2FC(Ctrl vs LPS)
Mean tags
S7B
IL4 H3K27ac CUT&Tag
upregulated 118
downregulated 117
Cd44
Snx29
Pde4b
Ptgs1
Mrc1
Pcyox11
Log2FC(Ctrl vs IL4)
Mean tags
S7C
GW3965 H3K27ac CUT&Tag
upregulated 126
downregulated 89
Scd1
Zfp871
Srebf1
Abcg1
Scd2
Lrch1
Log2FC(Ctrl vs GW3965)
Mean tags
S7D
IL4
159
61
3
12
2930
54
146
GW
LPS
S7E
LPS-regulated peaks
IL4-regulated peaks
GW-regulated peaks
Promoter
Enhancer
75.8%
75.3%
73.5%
Peak strength
Log10(abs(distance to TSS))
level
(0.0, 0.1]
(0.1, 0.2]
(0.2, 0.3]
(0.3, 0.4]
(0.4, 0.5]
(0.5, 0.6]
(0.6, 0.7]
(0.7, 0.8]
(0.8, 0.9]
(0.9, 1.0]
S7F
RAW264.7
shGFP
shNCOR
shSMRT
Tags counts (rpkm)
Stat6
Jun
Rela
S7G
Ccl2
Abca1
Mmp9
siCTRL
siNCOR
siSMRT
Relative mRNA levels
Ctrl
LPS
ns
*
**
***

**Supplementary Figure S7.** Analysis of NCOR vs. SMRT-dependent macrophage reprogramming in response to different signals.

(**A-C**) MA plots showing upregulated and downregulated H3K27ac peaks in response to LPS (**A**), IL4 (B) and GW3965 (**C**) (*n*=2). Representative upregulated peaks are highlighted with the annotated gene. (**D**) Venn diagram illustrating the overlap between altered H3K27ac peaks induced by LPS, IL4, and GW3965 in macrophages. (**E**) Distribution between promoters and enhancers of activated H3K27ac regions by LPS, IL4 and GW3965. The distribution is presented along the distance from the TSS of the annotated gene. (**F**) RNA-seq tag counts (-RPKM) showing gene expression levels of the TFs Stat6, Jun and p65 (Rela) in NCOR- vs. SMRT- depleted cells. (**G**) RT-qPCR analysis of related gene expression in LPS treatment in siNCOR and siSMRT RAW cells (*n*=4). Unpaired *t* test was used to determine data significance for gene expression. All data were represented as mean ± SEM. *$P < 0.05$, **$P < 0.01$, ***$P < 0.001$.

### Supplementary Table 1

#### PCR primers, shRNA oligos and siRNA oligos used in this study

| RT-qPCR primers | Forward Primer (5'-3') | Reverse Primer (5'-3') |
|---|---|---|
| *Gapdh* | ATGGCCTTCCGTGTTCCTA | TGAAGTCGCAGGAGACAACCT |
| *Ncor* | TGGATCCTGCTGCTGCTTACCT | GGCTGCTCTCGTGGGGACAGT |
| *Smrt* | GCCCTTAGTCCTAGGTGTGG | TTGTACAGAGGCGTGTGGGA |
| *Ptgs1* | CTTAGGCCACGGGGTAGA | TTTCCCATCCTTGAAGAGCCG |
| *Arg1* | CATTTGGGTGGATGCTCACA | TGGTACATCTGGGAACTTTCCTTT |
| *Abca1* | CCCAGAGCAAAAAGCGACTC | GGTCATCATCACTTTGGTCCTTG |
| *Abcg1* | CAAGACCCTTTTGAAAGGGATCTC | GCCAGAATATTCATGAGTGTGGAC |
| *Ccl2* | CAGATGCAGTTAACGCCCCA | TGAGCTTGGTGACAAAAACTACAG |
| *Pdcd1* | GAGCTCGTGGTAACAGAGAGAA | GAGTGTCGTCCTTGCTTCCA |
| *Ccl3* | AACCAAGTCTTCTCAGCGCC | GTCAGGAAAATGACACCTGGCTG |
| *Mmp9* | CAAGGACGGTTGGTACTGGA | GGATCCTCAAAGGCGGAGTC |
| *Chst1* | GATAGCGTCAGGAACGAGGG | ACACACAGCAGTTACCTTCCC |
| **shRNA oligos** | **Targets** | **Construct ID** |
| shGFP | GCAAGCTGACCCTGAAGTTCA | |
| shNCOR -1 | ATGGACAGGATCACGTATATT | TRCN0000350169 |
| shNCOR -2 | GTGATCATCACCCGGCAAATT | TRCN0000310755 |
| shSMRT-1 | TCCTCGCTGGCCCTCAATTAT | TRCN0000238140 |
| shSMRT-2 | TGTTGGCTTACAGGGTATATT | TRCN0000238139 |
| **siRNA oligos** | **Targets** | **Catalog ID** |
| siCTRL SMARTpool | UGGUUUACAUGUCGACUAA | D-001810-10-20, Dharmacon |
| | UGGUUUACAUGUUGUGUGA | |
| | UGGUUUACAUGUUUUCUGA | |
| | UGGUUUACAUGUUUUCCUA | |
| siNCOR SMARTpool | GCAGUGGAAGGAAGUAUAA | L-058556-00-0020, Dharmacon |
| | GAAAUCCCACGGCAAGAUA | |
| | CAACAACUCAGGUCAAUCA | |
| | CCAGGUCGAUGACAAGUGA | |
| siSMRT SMARTpool | GGCAAAGCCCACUGACUUA | L-045364-00-0020, Dharmacon |
| | GCAUGAGGUUUCUGAGAUC | |
| | GAAUGAGGUUCCCAGAGUU | |
| | GUACCCACCUUACCUCAUC | |

### Supplementary Table 2

Antibodies used in this study

| Antibody | Manufacturer | Item number | Application |
|---|---|---|---|
| NCOR | Bethyl | A301-145A | WB/ChIP-seq |
| SMRT | Bethyl | A301-147A | WB/ChIP-seq |
| GAPDH | Proteintech | 60004-1-Ig | WB |
| GPS2 | Homemade | N/A | WB |
| HDAC3 | Beyotime | AF2011 | WB |
| H3K27ac | Abcam | ab4729 | ChIP-seq/CUT&Tag |
| Jun | Santa Cruz | sc-166540 | WB |
| β-actin | Abcam | ab8226 | WB |
| Lamin B1 | Beyotime | AF1408 | WB |
| β-actin | FDbio | FD0060 | WB |
| STAT6 | Cell Signaling | 9362 | WB |
| p65 | Santa Cruz | sc-71675 | WB |

### Supplementary Table 3
Public datasets used in this study

| GSM number | Antibody | Type | Tissue | Treatment | GEO series |
|---|---|---|---|---|---|
| GSM1232940 | H4K5ac | ChIP-seq | Peritoneal macrophages | Basal | GSE50944 |
| GSM1232941 | H4K5ac | ChIP-seq | Peritoneal macrophages | Basal | GSE50944 |
| GSM4848548 | RUNX1 | ChIP-seq | RAW264.7 | Basal | GSE130383 |
| GSM4848549 | RUNX1 | ChIP-seq | RAW264.7 | Basal | GSE130383 |
| GSM4848528 | PU.1 | ChIP-seq | RAW264.7 | Basal | GSE130383 |
| GSM4848529 | PU.1 | ChIP-seq | RAW264.7 | Basal | GSE130383 |
| GSM4848572 | MED1 | ChIP-seq | RAW264.7 | Basal | GSE130383 |
| GSM4848573 | MED1 | ChIP-seq | RAW264.7 | Basal | GSE130383 |
| GSM4848609 | CBP | ChIP-seq | RAW264.7 | Basal | GSE130383 |
| GSM4848610 | CBP | ChIP-seq | RAW264.7 | Basal | GSE130383 |
| GSM4848584 | JunB | ChIP-seq | RAW264.7 | Basal | GSE130383 |
| GSM4848585 | JunB | ChIP-seq | RAW264.7 | Basal | GSE130383 |
| GSM4848503 | H3K27ac | ChIP-seq | RAW264.7 | shGFP rp1 | GSE130383 |
| GSM4848504 | H3K27ac | ChIP-seq | RAW264.7 | shGFP rp2 | GSE130383 |
| GSM4848507 | H3K27ac | ChIP-seq | RAW264.7 | shSMRT rp1 | GSE130383 |
| GSM4848508 | H3K27ac | ChIP-seq | RAW264.7 | shSMRT rp2 | GSE130383 |
| GSM5599368 | GPS2 | ChIP-seq | RAW264.7 | shGFP rp1 | GSE184884 |
| GSM5599369 | GPS2 | ChIP-seq | RAW264.7 | shGFP rp2 | GSE184884 |
| GSM5599372 | GPS2 | ChIP-seq | RAW264.7 | shNCOR rp1 | GSE184884 |
| GSM5599373 | GPS2 | ChIP-seq | RAW264.7 | shNCOR rp2 | GSE184884 |
